# PICALM deficiency exacerbates amyloid pathology in murine and nematode Alzheimer’s disease models

**DOI:** 10.64898/2026.11.23.768482

**Authors:** Domenico Azarnia Tehran, Gaga Kochlamazashvili, Christian Gallrein, Mira Sleiman, Sara Ayala Mariscal, Jasmeet Kaur Shergill, Kunal Gharat, Kinga Konkel, Max Ruwolt, Heike Stephanowitz, Mei Wu, Claudia Bahnik, Delia Löwe, Giorgia Carai, Fan Liu, Janine Kirstein, Volker Haucke, Tanja Maritzen

**Affiliations:** Department of Molecular Pharmacology and Cell Biology, Leibniz-Forschungsinstitut für Molekulare Pharmakologie (FMP), Robert-Rössle-Straße 10, 13125 Berlin, Germany; Department of Structural Interactomics, Leibniz-Forschungsinstitut für Molekulare Pharmakologie (FMP), Robert-Rössle-Straße 10, 13125 Berlin, Germany; Department of Biomedical Sciences, University of Padova, Via Ugo Bassi 58/B, 35131 Padua, Italy; Leibniz Institute on Aging – Fritz-Lipmann-Institute (FLI), Beutenbergstrasse 11, 07745 Jena, Germany; Department of Nanophysiology, RPTU University Kaiserslautern-Landau, Paul-Ehrlich-Straße 23, 67663 Kaiserslautern, Germany; Institute of Chemistry and Biochemistry, Freie Universität Berlin, Thielallee 63, 14195 Berlin, Germany; Friedrich-Schiller-Universität, Institute for Biochemistry & Biophysics, Hans Knöll-Strasse 2, 07745 Jena, Germany; Faculty of Biology, Chemistry, Pharmacy, Freie Universität Berlin, Takustraße 6, 14195 Berlin, Germany; Charité-Universitätsmedizin Berlin, NeuroCure Cluster of Excellence, Chariteplatz 1, 10117 Berlin, Germany

## Abstract

Alzheimer’s disease (AD), characterized by the accumulation of amyloid β-peptide (Aβ) plaques, is a progressive age-related neurodegenerative disorder affecting more than 35 million people worldwide. The risk of developing AD is significantly influenced by genetic factors. Deciphering their molecular functions has substantially advanced our understanding of the pathogenic mechanisms underlying AD. The PICALM gene represents a highly validated AD risk factor, and the encoded endocytic adaptor CALM is critical for endothelial Aβ clearance and microglial Aβ phagocytosis. However, it remains controversial how neuronally expressed CALM modulates AD pathogenesis. We find that loss of neuronal CALM accelerates Aβ plaque formation and impairs neurotransmission in an AD mouse model, revealing a protective role for neuronal CALM. These phenotypes are accompanied by an elevated surface level of the endolysosomal SNARE protein VAMP8, consistent with a role for CALM in promoting the turnover of aggregated proteins via the autophagy/lysosome pathway. Moreover, loss of CALM promotes Aβ aggregation in a *C. elegans* model expressing fluorescently labelled Aβ, while neuron-specific overexpression of CALM decreases Aβ aggregation. Finally, we demonstrate that the ANTH domain of CALM displays chaperone-like activity by inhibiting Aβ aggregation *in vitro*. Our combined data suggest that neuronal CALM might counteract AD progression by suppressing Aβ aggregation and by safeguarding endolysosomal SNARE trafficking.

## INTRODUCTION

Alzheimer’s disease (AD) remains the leading cause of dementia affecting approximately 33% of people older than 85 years and is therefore predicted to rise further with the aging population (Association, 2025). Two of its central hallmarks are the intracellular accumulation of hyperphosphorylated tau in neurofibrillary tangles (NFTs) and the aggregation of toxic amyloid β-peptides (Aβ) into extracellular plaques (Association, 2025). Aβ arises from the proteolytic processing of the amyloid precursor protein (APP). In the amyloidogenic pathway, APP is first cleaved by the β-secretase BACE1 and, subsequently, by Ψ-secretase, a multiprotein complex comprising presenilin and nicastrin (Wang et al., 2024). These cleavages occur mostly within intracellular compartments (Toh and Gleeson, 2016; Wang et al., 2024) and lead to the secretion of different Aβ species, including Aβ40 and Aβ42, with Aβ42 being particularly aggregation-prone and toxic in its oligomerized state. In addition to forming extracellular plaques, secreted Aβ can enter cells by dynamin-dependent endocytosis, thereby accumulating in the endolysosomal system (Ditaranto et al., 2001; Marshall et al., 2020; Schutzmann et al., 2021). Endolysosomal dysfunction, a characteristic of AD, can drive intraneuronal Aβ accumulation within lysosomes (Lee et al., 2022) which has been proposed to compromise their integrity and, thereby, function (Ditaranto et al., 2001; Lee et al., 2022; Liu et al., 2010; Zaretsky et al., 2022). Damaged lysosomes as well as constitutive neuron-specific endolysosomal perforations (Sanyal et al., 2025) likely allow Aβ access to the cytosol where it has been reported to inhibit proteasomal activity (Almeida et al., 2006; Oh et al., 2005; Tseng et al., 2008). In line with an important pathogenic function of intraneuronal Aβ, its removal in an AD mouse model at disease stages preceding extracellular plaque formation prevented early cognitive impairment (Billings et al., 2005).

While the mechanisms triggering late-onset AD (LOAD) remain incompletely understood, genome-wide association studies have uncovered multiple genetic risk loci, including variants of the PICALM gene (Bellenguez et al., 2022; Harold et al., 2009), which encodes the endocytic adaptor protein CALM. A harmful single nucleotide polymorphism in PICALM decreases CALM expression in endothelial cells (Zhao et al., 2015) and microglia (Kozlova et al., 2025); in the latter, this was traced to the reduced promoter binding of the transcription factor PU.1 (Kozlova et al., 2025). In endothelial cells, reduced CALM levels impair the uptake of Aβ via the clearance receptor LRP1, thereby decreasing the removal of Aβ across the blood brain barrier (Zhao et al., 2015). In microglia, reduced CALM levels cause an increase in lipid droplets and reactive oxygen species (ROS), possibly due to deficient lipid droplet degradation downstream of lysosomal dysfunction, resulting in impaired phagocytosis of Aβ and damaged myelin (Kozlova et al., 2025). In addition, the *Drosophila* CALM ortholog was shown to facilitate neuron-to-glia lipid transport and subsequent lipid droplet formation upon ROS induction (Moulton et al., 2021).

Although CALM, within the brain, is most abundant in endothelial cells and microglia, it is also expressed, albeit at lower levels, in neurons, where it fulfils important physiological functions. Super-resolution microscopy and fractionation experiments place CALM at neuronal synapses (Azarnia Tehran et al., 2022). At the presynapse CALM serves overlapping roles with its close relative AP180 in facilitating the endocytosis of the synaptic vesicle SNARE protein VAMP2 (Koo et al., 2015) as evidenced by VAMP2 accumulation on the surface of CALM-depleted neurons (Koo et al., 2011). At the postsynapse CALM is crucial for the internalization of Ca^2+^-permeable AMPA receptors, thereby modulating long-term depression and long-term potentiation (Azarnia Tehran et al., 2022).

However, the role of neuronal CALM in AD has remained highly controversial. CALM has been shown to associate with NFTs (Ando et al., 2013; Ando et al., 2016), and the reduction of CALM levels was reported to exacerbate tau pathology (Ando et al., 2020), but not tau spreading (Ando et al., 2026). Moreau *et al*. have proposed a protective effect of neuronal CALM by mediating the endocytosis of the SNARE proteins VAMP2, VAMP3 and VAMP8, thereby safeguarding autophagy and tau clearance (Moreau et al., 2014). Tian *et al*. suggested that AP2/PICALM, by interacting with the autophagy protein LC3 and the Aβ-amyloid precursor APP-CTF bridges Aβ degradation and autophagy (Tian et al., 2013). Treusch *et al*. suggested a beneficial function by demonstrating that CALM overexpression in cultured neurons ameliorates the toxic effects of exogenously applied Aβ oligomers (Treusch et al., 2011). Likewise, in a *Drosophila* model of Aβ42 toxicity, the CALM ortholog rescued Aβ toxicity which was proposed to occur via an effect on glutamatergic signaling (Yu et al., 2020). In contrast, studies using AD mouse models in conjunction with partial loss of CALM in heterozygous animals (Kanatsu et al., 2016) or via CALM-targeting shRNA (Xiao et al., 2012) have reported a decrease in Aβ plaque formation, suggesting that CALM exacerbates AD pathology. This was attributed to its facilitation of either nicastrin (Kanatsu et al., 2016; Kanatsu et al., 2014) or APP (Xiao et al., 2012) endocytosis, thereby promoting the amyloidogenic processing of APP. Taken together, these findings have led to conflicting models, in which neuronal CALM either limits proteotoxic stress by supporting endolysosomal homeostasis or enhances amyloidogenic APP trafficking and subsequent Aβ production.

In the present study we address these discrepancies with the aim of better understanding the potential contribution of neuronal CALM in the pathogenesis of AD. We have combined *in vitro* approaches with the generation of two animal models. First, we crossed APP transgenic mice with a CALM knockout (KO) line (Azarnia Tehran et al., 2022) that efficiently deletes CALM in excitatory forebrain neurons. Second, we deleted or neuronally overexpressed CALM in a *C. elegans* AD model that expresses fluorescently labelled Aβ42 in a pan-neuronal manner (Gallrein et al., 2021). Using these complementary *in vivo* models combined with *in vitro* experiments, we have tested different hypotheses regarding the function of neuronal CALM in AD. Our data support a protective role for neuronally expressed CALM that complements its established endothelial and microglial functions by counteracting Aβ aggregation.

## RESULTS

### Neuronal CALM deficiency aggravates AD pathology in mice

To study the effect of CALM loss on the progression of AD, we first crossed floxed CALM mice (CALM^lox^) with the EMX1-Cre mouse generated by Iwasato *et al*. that efficiently deletes in forebrain excitatory neurons (Iwasato et al., 2000). We previously demonstrated that the successful deletion of CALM in this mouse model impacts synaptic plasticity by controlling the internalization of Ca^2+^-permeable AMPA receptors from the neuronal surface (Azarnia Tehran et al., 2022). These CALM KO mice (CALM^E-ko^) (Azarnia Tehran et al., 2022) were further crossed into the established AD mouse model APP23 (Sturchler-Pierrat et al., 1997) (Supplementary Fig. 1A), which overexpresses the human APP splice isoform APP751 harboring the Swedish double mutation K670N/M671L. APP23 mice display clear Aβ deposition at one year of age and a concomitant decrease in basal synaptic transmission (Roder et al., 2003; Sturchler-Pierrat et al., 1997; Ye et al., 2017).

Our analysis revealed increased levels of soluble and insoluble Aβ40 and Aβ42 (Fig. 1A-D) in 18 month-old AD/CALM^E-ko^ mice, resulting in an overall higher Aβ burden (Fig. 1E) compared to AD mice expressing wild-type levels of neuronal CALM. Consistently, we found a significantly increased number and size of Aβ plaques (Fig. 1F-H) in 18 month-old mice, a phenotype that was accompanied by elevated GFAP levels (Supplementary Fig. 1B, C) indicative of reactive astrocytes and suggesting local neuroinflammation. Neuronal CALM deficiency thus aggravates AD pathology in mice.

**Fig. 1.**
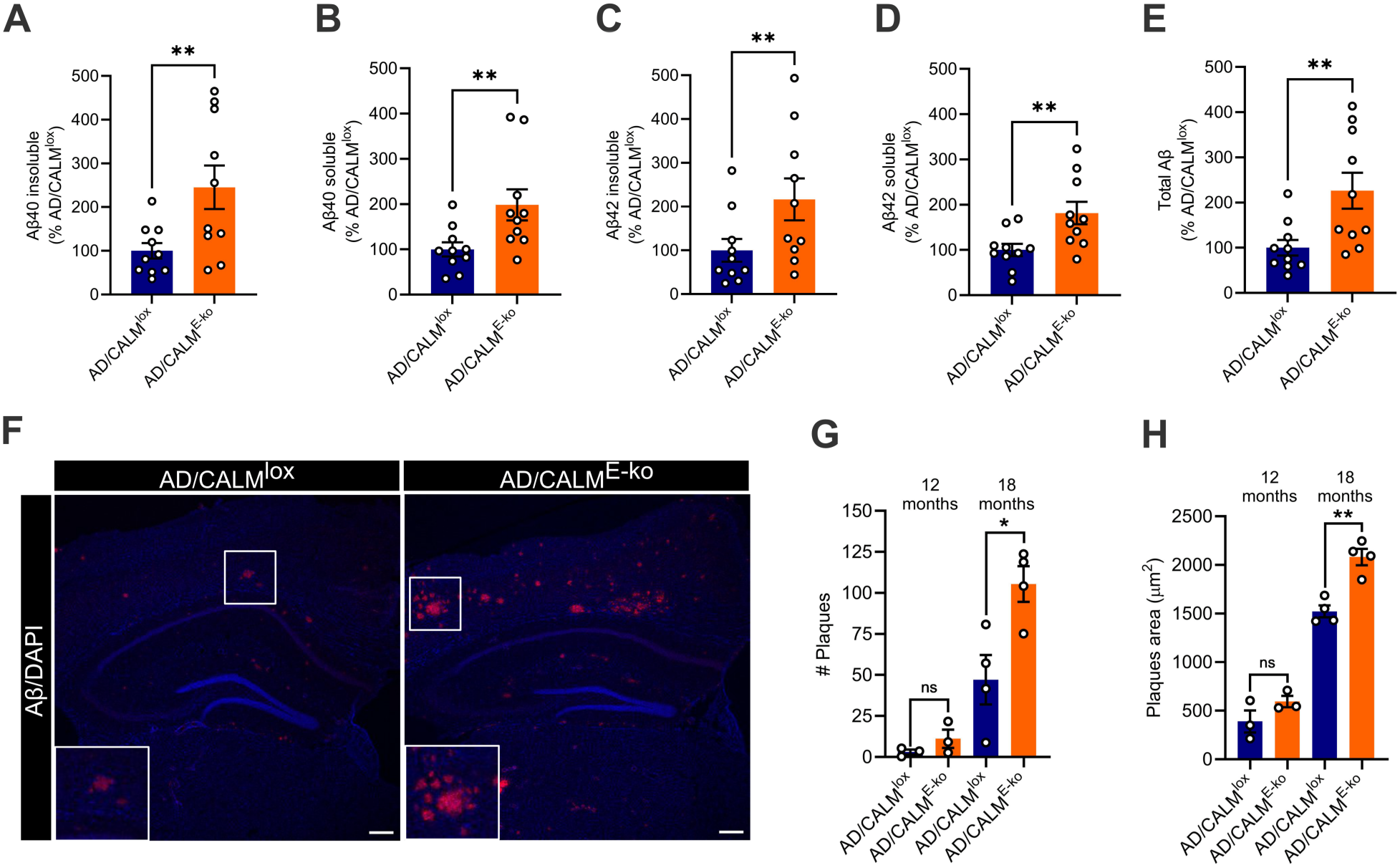
Neuronal CALM deficiency aggravates pathology in the Alzheimer’s disease (AD) mouse model APP23. (**A-E**) CALM loss increases Aβ load. Quantification of soluble and insoluble Aβ40 and Aβ42 as well as total Aβ by ELISA in homogenates derived from brains of 18-month-old female AD/CALM^lox^ and AD/CALM^E-ko^ mice. N=10. Two-tailed paired t test (A, D, E), Wilcoxon matched-pairs signed rank test (B, C). (**F-H**) CALM loss increases plaque formation. (**F**) Representative images of hippocampal brain sections of AD/CALM^lox^ and AD/CALM^E-ko^ mice stained with the Aβ-specific antibody 6E10 and DAPI to label nuclei. Scale bar, 200 µm. (**G-H**) Quantification of plaque number and area in 12- and 18-month-old female AD/CALM^lox^ and AD/CALM^E-ko^ mice. N=3-4. Two-tailed unpaired t test with Welch’s correction. For all quantifications data are depicted as mean±SEM. *=p<0.05, **=p<0.01, ns=non significant.

### Neuronal CALM loss impairs basal synaptic transmission in an AD model

To study potential effects of CALM loss on basal synaptic transmission in the AD mouse model, we recorded input/output stimulus response curves of field excitatory post-synaptic potentials (fEPSPs) in the CA1 region of the hippocampus, induced by stimulation of Schaffer collaterals/commissural fibers.

As previously described (Roder et al., 2003), mice carrying the APP23 transgene (AD/CALM^lox^) exhibited reduced fEPSPs already at 12 months of age compared to controls (CALM^lox^) (Fig. 2A). Neuronal CALM deficiency further aggravated the impairment of synaptic transmission in AD/CALM^E-ko^ mice (Fig. 2B), whereas basal neurotransmission remained unaffected in CALM^E-ko^ mice lacking the APP23 transgene (Fig. 2C). In all cases, fiber volley amplitudes, a readout for presynaptic excitability, were unchanged (Supplementary Fig. 1D-F). Finally, CALM deficiency also resulted in a significantly lower brain weight of AD/CALM^E-ko^ mice at 18 months of age (Supplementary Fig. 1G), while their overall body weight was unaltered (Supplementary Fig. 1H), further underlining the protective effect of CALM in the context of AD. These data demonstrate that neuronal CALM deficiency exacerbates synaptic dysfunction in an AD mouse model *in vivo*.

**Fig. 2.**
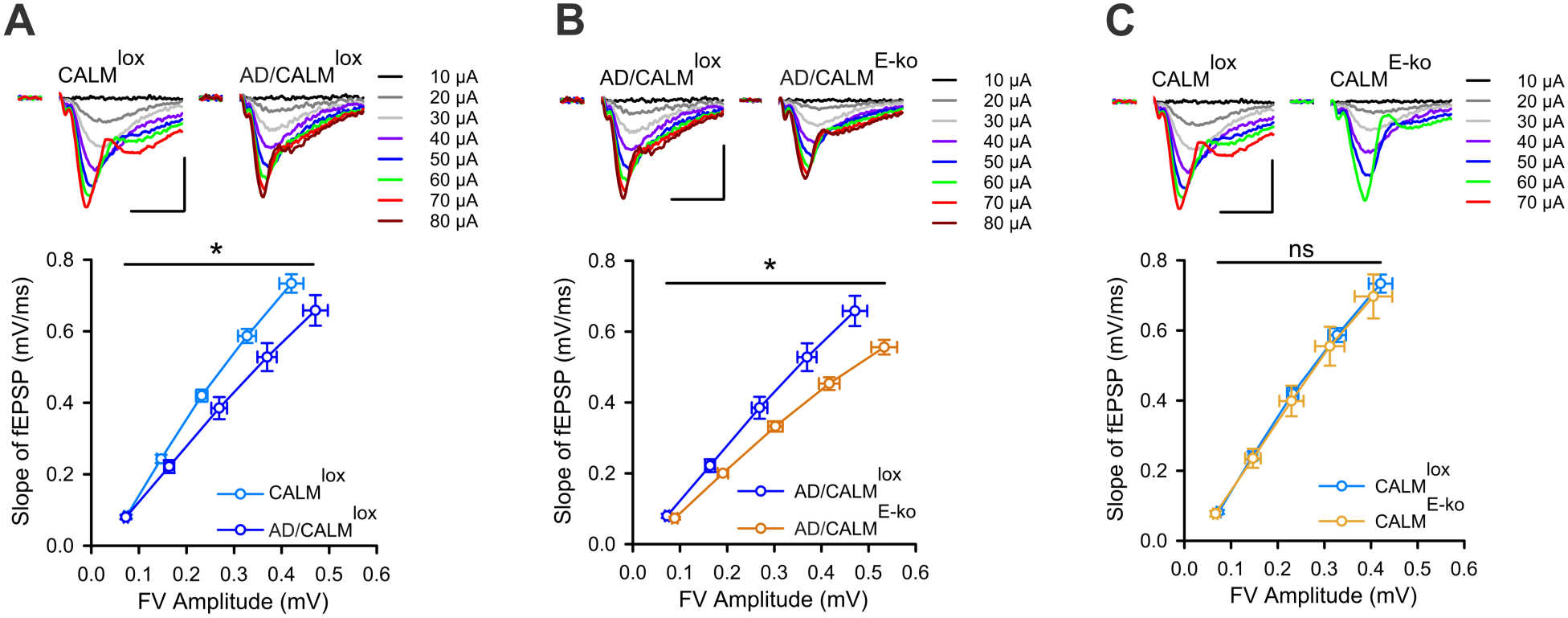
CALM loss impairs basal synaptic transmission in the Alzheimer’s disease mouse model APP23. (**A-C**) Top: Exemplary traces recorded in slices of the indicated genotypes evoked by increasing stimulation intensities until maximal fEPSPs were obtained. Scale bars, 1 mV and 10 ms. Bottom: The relationship between slopes and fiber volleys of fEPSPs over a range of stimulation intensities reveals significantly reduced basal neurotransmission in the AD mouse model (**A**) which is further aggravated by CALM loss (**B**) while CALM deficiency on its own does not impair neurotransmission of 12-month-old female mice (**C**). CALM^lox^: N=9 animals/n=21 slices, AD/CALM^lox^: N=8 animals/n=21 slices, AD/CALM^E-ko^: N=9 animals/n=24 slices, CALM^E-ko^: N=6 animals/n=16 slices. Two-way repeated measures ANOVA. For all quantifications data are depicted as mean±SEM. *=p<0.05, ns=non significant.

### Neuronal CALM loss does not cause alterations in APP, its processing enzymes or Aβ receptors

To dissect the molecular mechanism underlying the role of neuronal CALM in counteracting AD pathology, we tested different hypotheses proposed by earlier studies (Kanatsu et al., 2016; Kanatsu et al., 2014; Moreau et al., 2014; Zhao et al., 2015). We first determined the overall protein levels of APP, its processing enzymes BACE1, presenilin-2 and nicastrin as well as the Aβ clearance receptor LRP1 by immunoblotting of brain lysates of AD/CALM^E-ko^ mice. The total steady-state protein levels of BACE1, presenilin-2 and nicastrin were all unaltered in AD/CALM^E-ko^ mice (Fig. 3A, B). In contrast, GFAP levels were increased in AD/CALM^E-ko^ mice in line with our observation that GFAP accumulates around Aβ plaques of AD/CALM^E-ko^ mice (Fig. 3A, B; Supplementary Fig. 1B, C). Because the best-characterized AD-related function of CALM in endothelial cells is to facilitate LRP1-mediated Aβ uptake, we next checked whether CALM depletion might also affect neuronal LRP1 (Chen et al., 2022) surface levels. However, the surface biotinylation of cultured CALM-deficient neurons (CALM^cag-ko^) derived from a tamoxifen-inducible CALM KO line (Supplementary Fig. 2A) failed to reveal an accumulation of LRP1 at the plasma membrane (Fig. 3C-E). Unaltered neuronal LRP1 surface levels in CALM KO neurons suggest that LRP1 endocytosis in neurons is largely independent of CALM, in contrast to its endocytosis in endothelial cells. This is consistent with a similarly cell-type specific endocytic adaptor function of CALM with regard to transferrin receptor. CALM is essential for internalizing transferrin receptor in hematopoietic cells, leading to death from anemia in constitutive CALM KO mice (Ishikawa et al., 2015), but dispensable for transferrin receptor uptake in many cell lines (Ishikawa et al., 2015; Meyerholz et al., 2005) including neurons (Azarnia Tehran et al., 2022), likely due to cell-type specific functional overlap with other endocytic adaptors.

**Fig. 3.**
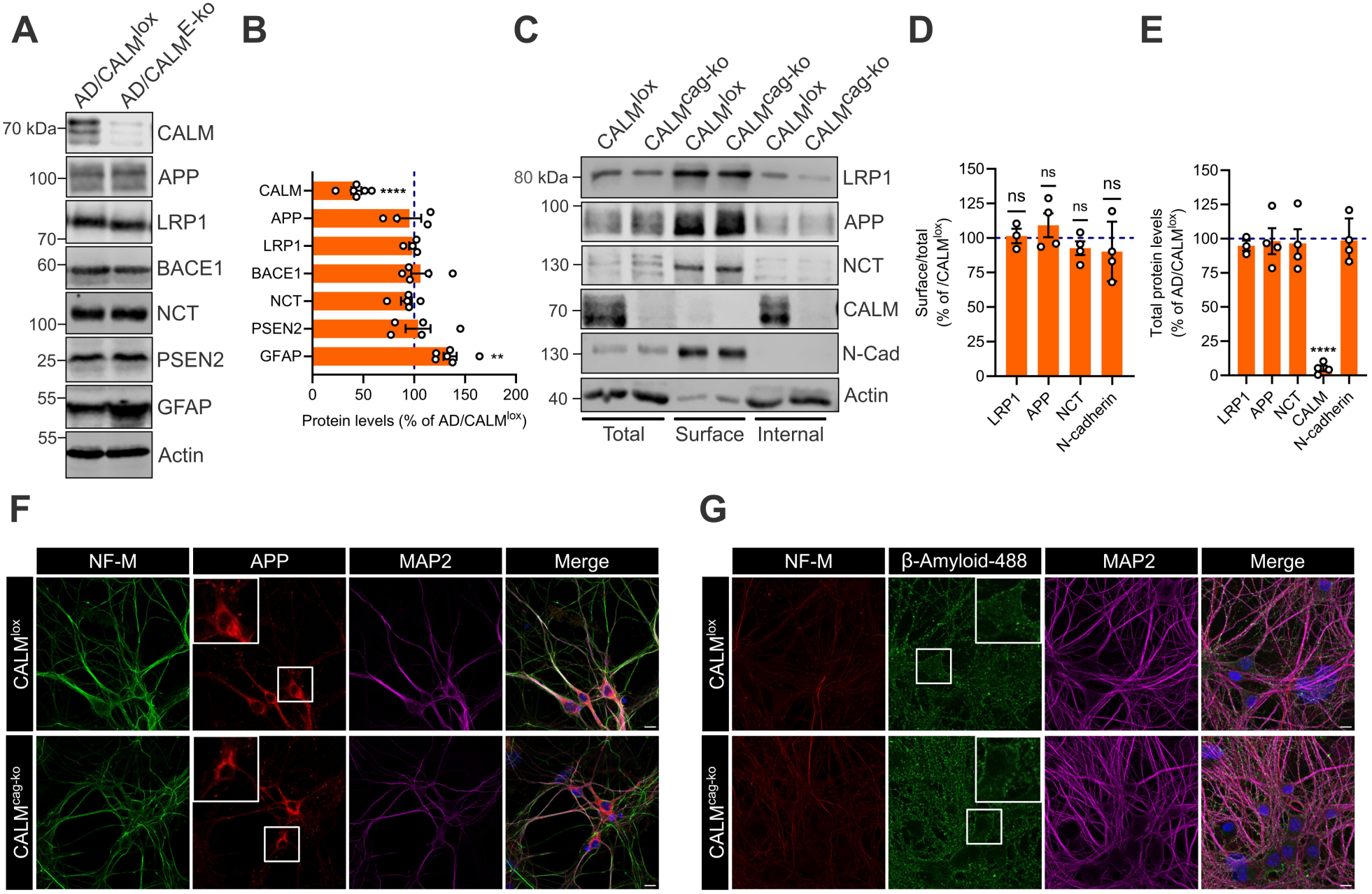
Neuronal CALM loss does not cause alterations in APP, the Aβ receptor LRP1, secretases or neuronal Aβ uptake. (**A, B**) Protein levels of APP, LRP1, β-secretase (BACE1) or the Ψ-secretase subunits nicastrin (NCT) and presenilin-2 (PSEN2) are unaltered upon CALM loss, while GFAP is increased. Total brain lysates from 18-month-old female AD/CALM^lox^ and AD/CALM^E-ko^ mice were probed by immunoblotting with the indicated antibodies. Representative immunoblot (**A**) and quantification (**B**) of protein levels in AD/CALM^E-ko^ brains normalized to AD/CALM^lox^ values. Dots represent individual animals, N=4-6. One sample t test. (**C-E**) CALM deletion does not alter surface levels of LRP1, APP or nicastrin (NCT). After surface biotinylation of cultured neurons derived from CALM^lox^ or CALM^cag-ko^ mice, total proteins, biotinylated surface proteins and intracellular proteins were analyzed by immunoblotting with the indicated antibodies. N-Cadherin (N-Cad) served as positive control and actin as negative control. (**C**) Representative immunoblots. (**D**) Surface/total ratios normalized to control. (**E**) Total levels normalized to control. (D, E) Dots represent independent neuronal cultures, N=3-4. One sample t test. (**F**) No change in APP localization. Cultured CALM^lox^ or CALM^cag-ko^ hippocampal neurons were fixed and stained with the indicated antibodies. (**G**) No change in neuronal Aβ uptake. Live cultured CALM^lox^ or CALM^cag-ko^ neurons were incubated at 37°C with fluorescently labelled Aβ to monitor its endocytosis. (F, G) Representative images from one independent experiment. Scale bar: 10 µm. NF-M, Neurofilament M. For all quantifications data are depicted as mean±SEM. **=p<0.01, ****=p<0.0001, ns=non significant.

Using the same surface biotinylation approach, we next assessed the surface levels of APP and nicastrin in CALM-deficient neurons. Plasma membrane levels of both proteins were likewise unaltered, suggesting that CALM does not control their steady-state surface distribution (Fig. 3C-E). Consistent with these findings, our immunofluorescence analysis of hippocampal neurons revealed no detectable alterations in the subcellular APP localization upon CALM loss (Fig. 3F). Finally, in agreement with the unaltered LRP1 surface levels, the uptake of fluorescently labelled Aβ42 by cultured neurons was unchanged upon neuronal CALM loss (Fig. 3G).

We conclude that the protective function of neuronal CALM with respect to Aβ42 plaque formation is likely independent of APP, LRP1 or Ψ-secretase endocytosis since a deficient internalization of these cargoes upon CALM deletion should have been reflected in increased surface levels, although minor functional defects cannot be ruled out. Based on these results, it also appears unlikely that neuronal CALM is required to preserve the general integrity of the endocytic pathway in neurons, unlike what has previously been observed in yeast (Treusch et al., 2011).

### Neuronal CALM sorts the endolysosomal SNARE protein VAMP8

Given the negative results reported above, we turned to a TMT-based mass spectrometry analysis of synaptosomes from AD/CALM^E-ko^ mice to address the mechanistic underpinnings of CALM’s protective effect in AD in an unbiased manner (Fig. 4A). This analysis revealed surprisingly few alterations in synaptic protein abundance. The most prominent change was a striking increase in VAMP8 and its cognate SNARE protein syntaxin7 (Mullock et al., 2000) in synaptosomes from AD/CALM^E-ko^ mice (Fig. 4B), as further confirmed by immunoblotting (Fig. 4C). These data are consistent with the known function of CALM as an endocytic adaptor for endolysosomal R-SNARE proteins of the VAMP family (Koo et al., 2015; Miller et al., 2011), which has been suggested to affect AD pathology by its effect on the sorting of VAMP2, VAMP3 and/or VAMP8 (Moreau et al., 2014). The observed elevation of total VAMP8 levels in synaptosomes likely originates from impaired targeting of VAMP8 from the cell surface to the endolysosomal system via CALM-mediated endocytosis (Koo et al., 2015; Miller et al., 2011). If this was true, the steady-state neuronal surface levels of VAMP8 should be elevated in the absence of CALM. Since VAMP8 does not contain any extracellular domain amenable to biotinylation, we employed a VAMP8 variant that is luminally fused to the pH-sensitive fluorescent protein pHluorin in conjunction with an established acid-base quench protocol (Koo et al., 2011) to quantify VAMP8 surface levels. This analysis revealed a prominent surface accumulation of VAMP8 in CALM-deficient neurons (Fig. 4D,E), in agreement with earlier data demonstrating that CALM binds VAMP8 and facilitates its internalization in HeLa cells (Miller et al., 2011). In summary, these data suggest that CALM facilitates the endocytic removal of VAMP8 from the plasma membrane also in neurons *in vivo*. We therefore hypothesize that the missorting of VAMP8 in absence of CALM could affect VAMP8/syntaxin7-dependent membrane fusion within the autophagy/lysosome pathway which has been implicated in the clearance of Aβ aggregates (Nixon et al., 2005; Pickford et al., 2008; Spilman et al., 2010). Whether this is sufficient to account for the increased Aβ burden of AD/CALM^E-ko^ mice remains to be directly tested.

**Fig. 4.**
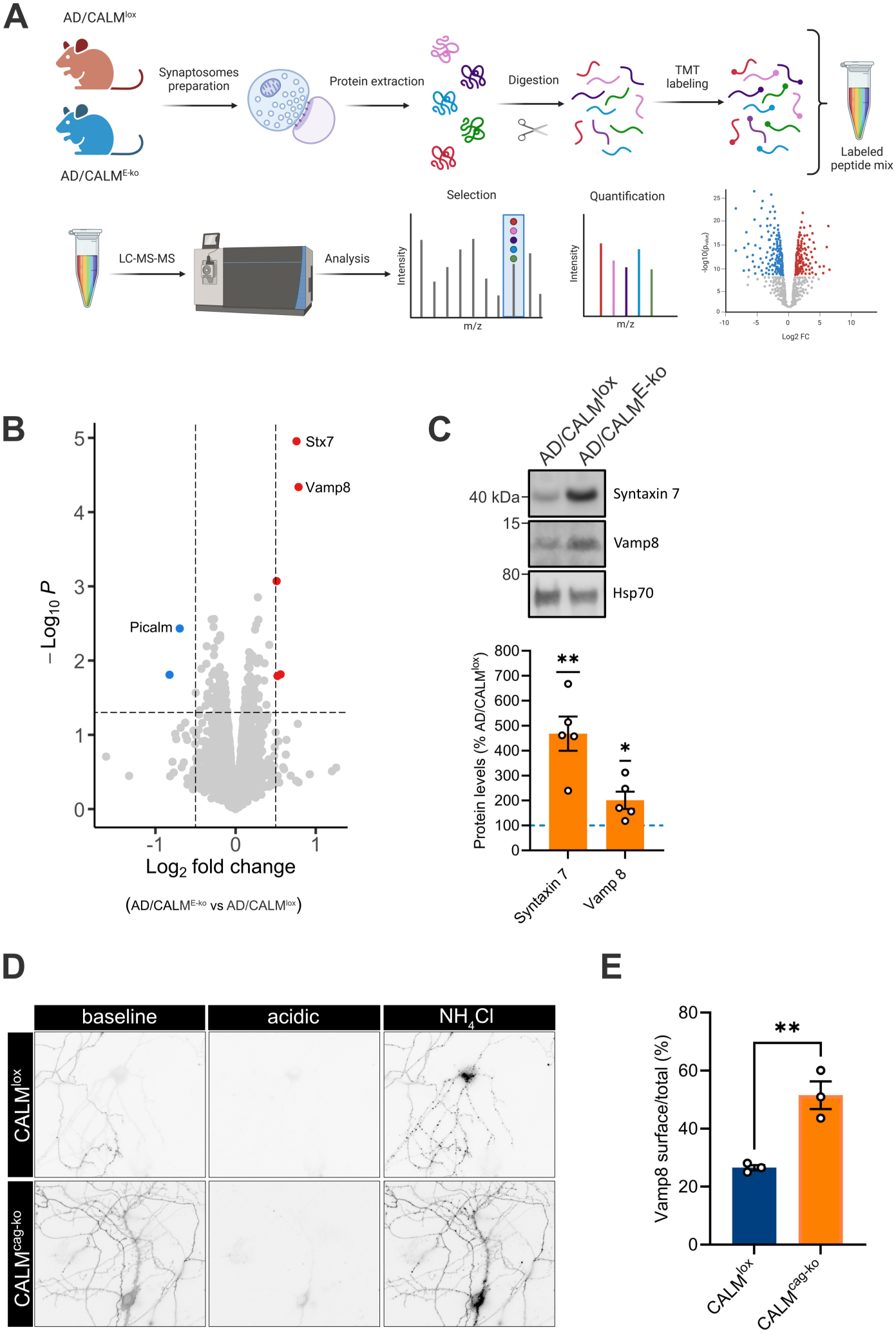
CALM selectively alters the abundance of the endolysosomal SNARE proteins VAMP8 and syntaxin7. (**A-C**) Loss of CALM leads to an increase in VAMP8 and syntaxin7 at synapses of AD/CALM^E-ko^ mice. (**A**) Scheme of proteomics experiment. (**B**) Volcano plot of mass spectrometry-derived synaptic proteome of AD/CALM^lox^ vs AD/CALM^E-ko^ mice. (**C**) Western blot-based confirmation of elevated VAMP8 and syntaxin7 levels. Bottom: Quantification of protein levels in AD/CALM^E-ko^ mice normalized to AD/CALM^lox^ levels. N=5 animals. One sample t test. (**D, E**) Surface accumulation of VAMP8 upon CALM loss. VAMP8-pHluorin-transfected hippocampal neurons derived from CALM^lox^ or CALM^cag-ko^ brains were probed for VAMP8 surface levels by acid-base quenching. (**D**) Representative images of VAMP8-pHluorin signals. (**E**) Quantification of VAMP8 surface fraction. N=3 independent experiments with neuronal cultures. Two-tailed unpaired t test. For all quantifications data are depicted as mean±SEM. *=p<0.05, **=p<0.01.

### CALM suppresses Aβ aggregation in living *C. elegans*

An influence of CALM on VAMP8 endocytosis is consistent with CALM’s canonical function as an endocytic sorting adaptor (Miller et al., 2011) and could contribute to the autophagic removal of Aβ aggregates. Still, two yeast studies motivated us to also explore the more unconventional hypothesis that CALM might affect Aβ aggregation. The first study established a yeast model expressing Aβ42 fused to an ER targeting signal, in which the secreted and subsequently endocytosed Aβ decreased cell growth. When screening for genetic modifiers of Aβ toxicity by overexpressing 5532 open reading frames, this study identified CALM as one of 23 suppressors of Aβ toxicity (Treusch et al., 2011). This was followed up by another yeast screen where Aβ was fused to a translational release factor the function of which was suppressed upon Aβ oligomerization. When testing candidates from the first screen, only the overexpression of the yeast CALM ortholog kept the release factor functional, suggesting that CALM harbors the propensity to suppress Aβ oligomerization (Park et al., 2016).

We therefore hypothesized that CALM, in addition to its endocytic function, may affect AD by serving as a chaperone that suppresses Aβ aggregation directly. To test this hypothesis, we first capitalized on an improved *C. elegans* AD model that sub-stoichiometrically expresses mScarlet-labelled Aβ42 to allow the detection of Aβ by fluorescence lifetime microscopy (FLIM) without perturbing fibril formation (Gallrein et al., 2021) (Supplementary Fig. 2B). Since the incorporation of mScarlet-Aβ into a growing amyloid fibril will lead to the quenching of the fluorophore and, thereby, to a reduction of the apparent fluorescence lifetime, FLIM allows Aβ aggregation to be quantified using the decrease of the fluorescence lifetime as a proxy (Gallrein et al., 2021). Deletion of the *C. elegans* PICALM homolog *unc-11* in nematodes expressing mScarlet-Aβ in neurons (Δ*unc-11* x nAβ), led to mildly increased Aβ aggregation as reflected by the reduction in fluorescence lifetime (Fig. 5A).

**Fig. 5.**
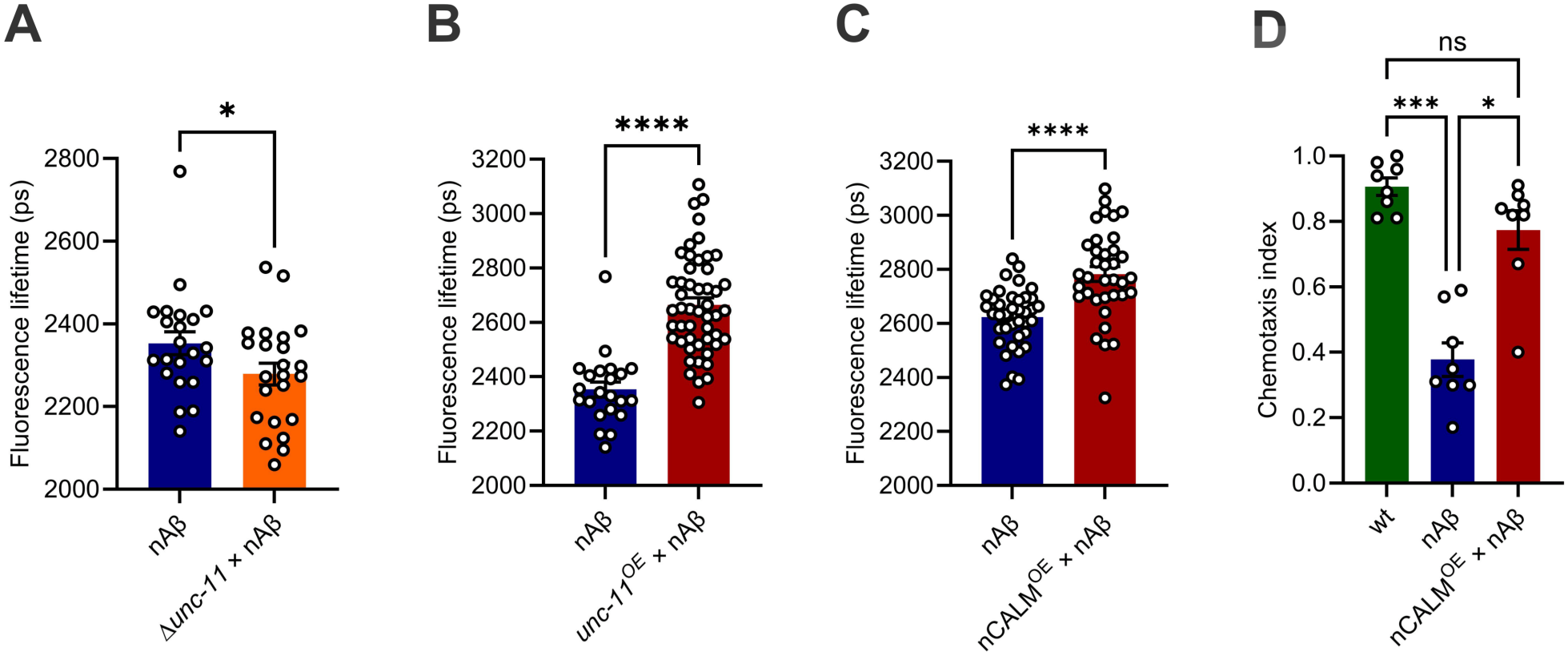
CALM suppresses Aβ aggregation in living *C. elegans*. (**A-C**) CALM suppresses Aβ aggregation in living nematodes. 7-day-old *C. elegans* neuronally expressing mScarlet-Aβ (nAβ) in combination with an unc-11 knockout phenotype (Δunc-11; **A**), unc-11 overexpression (unc-11OE; **B**) or neuronal overexpression of human CALM (nCALM^OE^; **C**) were subjected to fluorescence lifetime imaging. Mann-Whitney test (A, B), unpaired t test with Welch’s correction (C). Circles indicate analyzed animals ((A) N(nAβ) = 22, N(Δunc-11 x nAβ) = 23; (B) N(nAβ) = 22, N(unc-11^OE^ x nAβ) = 50; (C) N(nAβ) = 38, N(nCALM^OE^ x nAβ) = 36. (**D**) CALM overexpression protects *C. elegans* neuronally expressing mScarlet-Aβ from impairment of chemotactic behaviour. Quantification of chemotactic index of 7-day-old wild-type nematodes and mScarlet-Aβ expressing nematodes in absence or presence of neuronal CALM overexpression. N = 8 independent experiments with cohorts of 25-100 animals. Kruskal-Wallis test with Dunn’s multiple comparison test. For all quantifications data are depicted as mean±SEM. *=p<0.05, ***=p<0.001, ****=p<0.0001, ns=non significant.

Since the deletion of *unc-11* in *C. elegans* was not specific to neurons, we overexpressed *unc-11* specifically in neurons via a pan-neuronal promoter (Supplementary Fig. 2C) to test whether the increased Aβ aggregation upon *unc-11* deletion is due to neuronal CALM function. Neuronal overexpression of *unc-11* significantly decreased Aβ aggregation, as reflected by the substantial increase in fluorescence lifetime (Fig. 5B). Notably, neuronal expression of human CALM recapitulated this protective effect (Fig. 5C; Supplementary Fig. 2D).

To probe whether the decreased Aβ aggregation correlates with improved neuronal function, we performed chemotaxis assays. Expression of Aβ in *C. elegans* severely impairs their ability to respond by chemotaxis to a volatile compound, 2% benzaldehyde (Fig. 5D). The impaired chemotaxis was significantly restored by overexpression of human CALM (Fig. 5D), further supporting a protective role of CALM *in vivo*. These results are consistent with an earlier study employing a *C. elegans* model where Aβ is targeted to the secretory pathway of the five glutamatergic neurons present in the worm, resulting in 75% of nematodes losing one or more of these neurons by day 7. Overexpression of *unc-11* in this model increased the proportion of worms with five surviving neurons (Treusch et al., 2011).

Hence, neuronal CALM counteracts amyloid pathology also in *C. elegans in vivo* and appears to do so by affecting Aβ aggregation itself as demonstrated by the FLIM measurements.

### CALM suppresses Aβ aggregation *in vitro*

These data lend support to a model where neuronal CALM in addition to its conserved role in the endocytosis of SNARE proteins such as VAMP8 (Fig. 4), which have been implicated in the degradation of Aβ aggregates via the autophagy/lysosome pathway (Furuta et al., 2010; Moreau et al., 2014), might fulfill a further mechanistic role by acting as a molecular chaperone to directly suppress Aβ aggregation. To probe whether CALM has a direct effect on Aβ aggregation, we turned to an *in vitro* approach using purified proteins. CALM consists of an N-terminal ANTH domain that binds to SNARE proteins and the lipid PI(4,5)P_2_, and an intrinsi-cally disordered C-terminal tail containing motifs for the interaction with endocytic proteins (Azarnia Tehran et al., 2019). We used the ANTH domain of CALM fused to a 6xHis-tag in an Aβ aggregation assay where purified monomeric Aβ42 is incubated in the presence of the amyloid-specific fluorescent dye Thioflavin T, which allows Aβ aggregation to be quantified by the associated increase in Thioflavin T fluorescence (Biancalana and Koide, 2010). Addition of 1 µM or 5 µM CALM-ANTH domain to 5 µM Aβ strikingly suppressed Aβ aggregation, whereas addition of His-Δ51-Gadkin, an unrelated protein with functions in endosomal motility (Maritzen et al., 2010), which served as easy-to-purify protein as specificity control, had no impact (Fig. 6A,B), underlining that the observed chaperone-like effect is specific for CALM’s ANTH domain.

**Fig. 6.**
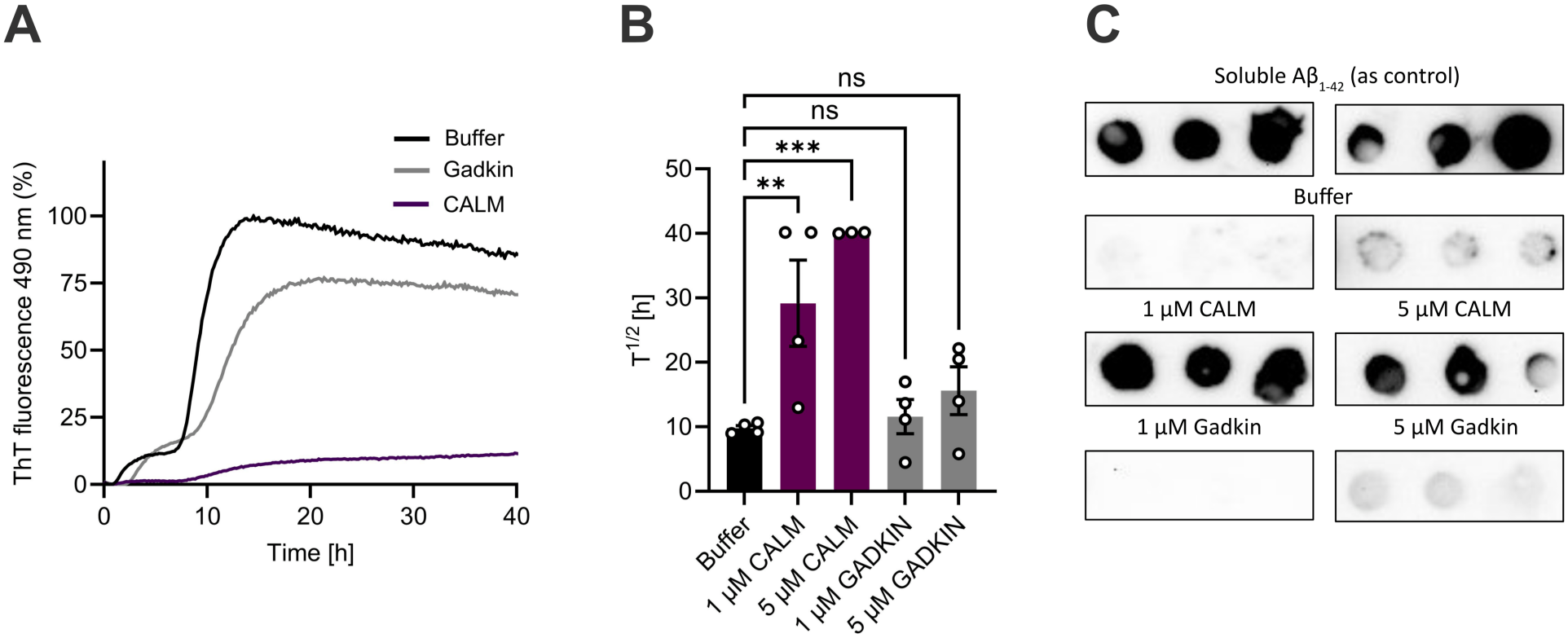
CALM’s ANTH domain suppresses Aβ aggregation *in vitro*. (**A**) Representative time-course plot of Thioflavin T fluorescence as proxy for the aggregation of 5 µM Aβ in presence of only buffer, 1 µM human His-CALM-ANTH domain or 1 µM His-β51-Gadkin as negative control. (**B**) Quantification of half-life of Aβ aggregation. Circles indicate independent experiments, N=3-4. One-way ANOVA with Holm-Sidak’s multiple comparison test. (**C**) Dot blot of initially monomeric Aβ and the remaining soluble Aβ fraction after 40 h aggregation in presence of buffer or the indicated amounts of human His-CALM-ANTH or His-β51-Gadkin. For all quantifications data are depicted as mean±SEM. **=p<0.01, ***=p<0.001, ns=non significant.

The suppression of Aβ aggregation by CALM-ANTH was validated by an analysis of the remaining soluble Aβ fraction via immunoblotting. For that, the supernatants of the same samples were spotted onto a membrane and immunostained with Aβ-specific antibodies. Monomeric Aβ served as control (Fig. 6C). Due to its efficient aggregation, soluble Aβ was hardly detectable after 40 h in samples containing only buffer or in the presence of His-Δ51-Gadkin. However, it was present at similar levels as the monomeric Aβ control after incubation with the CALM-ANTH domain (Fig. 6C). We conclude that CALM via its ANTH domain is sufficient to directly suppress Aβ aggregation while preserving soluble Aβ in a cell-free system with purified components, consistent with a chaperone-like activity (Klaips et al., 2018).

## DISCUSSION

Our data unravel that neuronal CALM, encoded by the established AD risk gene PICALM, significantly impacts AD pathology *in vivo*, potentially adding to the protective effects reported for endothelial (Zhao et al., 2015) and microglial (Kozlova et al., 2025) CALM. We find that loss of neuronal CALM aggravates Aβ plaque formation and impairs neurotransmission in APP transgenic mice, while the neuron-specific overexpression of CALM suppresses Aβ aggregation and proteotoxicity in Aβ-expressing nematodes.

Even though it remains uncertain exactly how neuronal CALM exhibits its protective effect with respect to AD pathology, our data support a subset of the previously proposed hypotheses. First, we find no evidence in support of the previously suggested roles of CALM in safeguarding neuronal APP or nicastrin internalization (see Fig. 3). In addition, in neurons CALM does not appear to act as a major endocytic adaptor for LRP1 (Fig. 3), in contrast to its important role in LRP1 uptake in endothelial cells (Zhao et al., 2015). Instead, in an unbiased mass spectrometry-based screen we observe a highly specific and profound effect of neuronal CALM loss on the overall protein amount as well as surface level of the endolysosomal SNARE protein VAMP8 at forebrain synapses from AD/CALM^E-ko^ mice (Fig. 4). This is consistent with earlier data demonstrating that CALM directly binds VAMP8 and facilitates its internalization in HeLa cells (Miller et al., 2011). Mutating the VAMP8 binding site in CALM results in the accumulation of VAMP8 at the plasma membrane of HeLa cells (Miller et al., 2011), similarly to what we observe in neurons upon CALM deletion.

Missorting of endolysosomal SNARE proteins including VAMP8 due to CALM depletion has previously been found to impair autophagy-dependent protein degradation (Furuta et al., 2010) and to exacerbate tau-mediated toxicity in a zebrafish transgenic model (Moreau et al., 2014). Our data support this hypothesis and establish a physiological relevance of neuronal CALM in endolysosomal SNARE sorting *in vivo*. Whether this phenotype underlies the aggravated Aβ plaque formation in our AD/CALM^E-ko^ mice remains to be directly tested, but a contribution seems plausible.

Finally, we find that CALM also displays a strong chaperone-like activity in ameliorating Aβ aggregation in *C. elegans in vivo* and in the test tube *in vitro*. It is therefore conceivable that neuronal CALM exerts a dual protective role in AD by inhibiting Aβ aggregation while at the same time promoting Aβ turnover via the autophagy/lysosome pathway.

Our observation that CALM suppresses the aggregation of Aβ *in vivo* and also in a controlled *in vitro* setting raises the question of where this interaction may occur *in vivo*. To our knowledge, CALM is a cytosolic protein, while Aβ is predominantly secreted into the extracellular space upon APP processing by Ψ-secretase (Zhang et al., 2011). We therefore envision a model in which extracellular Aβ is first endocytosed (Gallego Villarejo et al., 2022) (a step that is e.g. well documented in the context of endothelial Aβ clearance by transcytosis), followed by partial leakage from the lumen of endosomes and/or lysosomes into the cytosol. Several lines of evidence suggest that endocytosed Aβ can indeed undergo leakage into the cytoplasm. First, Aβ has been shown to exert detrimental effects on cytosolic protein complexes like the proteasome (Almeida et al., 2006; Oh et al., 2005; Tseng et al., 2008) and to gain access to mitochondria (Hansson Petersen et al., 2008). Second, a fraction of neuronal lysosomes has been demonstrated to be perforated (Sanyal et al., 2025) and, thereby to allow the cytosolic entry of amyloid proteins such as α-synuclein. Third, Aβ taken up into the endolysosomal system has been demonstrated to compromise lysosomal membrane integrity, thereby generating its own entry route into the cytosol (Ditaranto et al., 2001; Lee et al., 2022; Liu et al., 2010; Zaretsky et al., 2022). In support of this hypothesis, it has been observed that neurons with early intracellular Aβ accumulation are particularly vulnerable to cell death (Caramello et al., 2025).

Our data are thus consistent with (even though they do not yet prove) a mechanism, whereby CALM via its ANTH domain chaperones the escaped Aβ to prevent the formation of toxic Aβ assemblies within the cytosol. Suppressing fibril formation might at the same time facilitate the removal of cytosolic Aβ by mechanisms such as autophagy. With regard to this step, CALM might execute its second protective function by safeguarding the distribution of endolysosomal SNARE proteins which are critical for promoting autophagosome degradation via autophagosome fusion with lysosomes. Ultimately, the efficient disposal of intracellular Aβ may also help limiting extracellular Aβ plaque formation since toxic intracellular Aβ aggregates could contribute to extracellular plaques upon neuronal cell death and leakage into the extracellular space. In summary, our data are consistent with a dual role of CALM in neurons: the direct suppression of Aβ aggregation and the indirect promotion of aggregate removal by safeguarding SNARE trafficking. Enhancing CALM function across the different brain cell types may thus represent a strategy to counteract AD-related pathology.

### Limitations of the study

We note that the EMX1-Cre mouse line was shown in some studies to mediate recombination not only in excitatory forebrain neurons (Guo et al., 2000; Iwasato et al., 2000), but also in a population of neocortical astrocytes (Gorski et al., 2002) while not being active in endothelial cells (Gorski et al., 2002). Therefore, an astrocytic contribution to the *in vivo* phenotypes cannot be fully excluded. However, the convergence of our mouse model findings with the neuron-specific *unc-11*/CALM manipulations in *C. elegans* supports the loss of neuronal CALM as a major driver of the observed pathology. Clearly, further studies are needed to dissect the exact physiological role of neuronal CALM in counteracting Aβ and tau pathology *in vivo*, to establish whether VAMP8/syntaxin7 missorting is causally linked to the Aβ phenotype, and to dissect the relative contributions of neuronal, astrocytic, endothelial and microglial CALM in ameliorating AD progression.

## Materials and methods

### Mice

Mice were housed under standard conditions and monitored daily by trained animal caretakers. The animal facility was regularly checked for standard pathogens. Only female mice were used for this study. The age of each animal used is reported in the respective experimental sections. Mice were kept in groups of up to six animals in standard individually ventilated cages (IVC) of 524 cm^2^ containing bedding and nesting material. Food and water were provided ad libitum. The light cycle ran from 6 a.m. to 6 p.m. Forebrain-specific PICALM KO and tamoxifen-inducible PICALM KO mice are described in (Azarnia Tehran et al., 2022).The APP transgenic mouse strain Tg (Thy1-APPKM670/671NL)23 (commonly denoted APP23) is described in (Sturchler-Pierrat et al., 1997) and was heterozygously crossed into the two different PICALM mouse models. Throughout the text, PICALM^lox/lox^ Cre-negative mice without APP23 transgene are denoted as “CALM^lox^”, PICALM^lox/lox^ EMX1-Cre-positive mice without APP23 transgene are denoted as “CALM^E-ko^”, PICALM^lox/lox^ iCAG-Cre-positive mice without APP23 transgene are denoted as “CALM^cag-ko^”, PICALM^lox/lox^ Cre-negative mice carrying the APP23 transgene are denoted as “AD/CALM^lox^” and PICALM^lox/lox^ EMX1-Cre-positive mice with APP23 transgene are denoted as “AD/CALM^E-ko^”.

### Genotyping

For genotyping, genomic DNA was extracted from mouse tissue biopsies and analyzed with a standard PCR program using the primers indicated in Supplementary Table 1.

### Aβ measurements

Cortical and hippocampal tissues were collected from 18 month-old female mice and homogenized in ice-cold RIPA buffer (150 mM NaCl, 1% NP-40, 0.5% sodium deoxycholate, 0.1% SDS, 50 mM Tris-HCl), containing protease inhibitors (cOmplete™, Mini protease inhibitor cocktail, Merck) and phosphatase inhibitors (phosphatase inhibitor cocktail 2 and 3, Merck) using 20 strokes at 900 rpm. Protein homogenates were incubated for 30 min at 4°C followed by centrifugation at 1,000 g for 10 min to remove debris and myelin. The supernatants were centrifuged at 100,000 g for 1 h at 4°C. The supernatant, representing the RIPA-soluble fraction, was collected and stored at -20°C. The insoluble pellet was dissolved in formic acid and neutralized with 1 M Tris Base containing 0.1 M NaCl (pH 11) and stored at -20°C. Aβ40 and Aβ42 levels were measured using ELISA kits (Human/Rat β Amyloid(40) or β Amyloid(42) ELISA Kit Wako II) according to the manufacturer’s instructions.

### Immunohistochemistry

Mice were anesthetized by i.p. injection of ketamin/xylazin and transcardially perfused with PBS, followed by 4% (wt/vol) PFA in phosphate buffer (PB: 0.125 mM Na_2_HPO_4_/NaH_2_PO_4_, pH 7.4) at a flow rate of 8 ml/min. Brains were carefully taken out of the skull, postfixed overnight in the same fixative and placed in DMSO solution [a mixture of 20% (vol/vol) glycerol and 2% (vol/vol) dimethyl sulfoxide (DMSO) in 0.4 M PB] for 24 h for cryoprotection. Coronal sections (25 μm thick) were collected in six series in DMSO solution. For immunostaining, sections were washed for 3 h in PB (with solution exchange every 20 min), permeabilized for 2 h in PB containing 0.3% Triton-X-100, blocked for 1 h in 5% (vol/vol) normal goat serum and 0.3% Triton-X-100, and incubated 2-3 days at 4°C with primary antibodies (Supplementary Table 2). Unbound antibodies were removed by washing in PB for 2 h (with solution exchange every 15 min), and bound antibodies were decorated with Alexa Fluor-conjugated secondary antibodies (Supplementary Table 2) for 2 h in PB. Finally, nuclei were stained with DAPI (1 µg/ml in PB), and sections were mounted on gelatine-coated glass slides. Images of brain slices were acquired blinded to genotype using a Zeiss LSM 710 confocal laser scanning microscope with a 20x or a 40x objective. All acquisition settings were set equally for sections of all groups within each immunostaining.

### Electrophysiology

12 month-old female mice of all genotypes were used for acute hippocampal field electrophysiology. Mice were decapitated after cervical dislocation, and the brains quickly extracted into ice-cold dissection artificial cerebrospinal fluid (ACSF) containing (in mM): 2.5 KCl, 1.25 NaH_2_PO_4_, 24 NaHCO_3_, 1.5 MgS0_4_, 2 CaCl_2_, 25 glucose, 250 sucrose. The brains were dissected into equal hemispheres, glued on an object mounting platform and sectioned into 350 µm thick sagittal slices using vibroslicer (Leica, VT 1200S). The slices were collected in a resting chamber containing ACSF, in which 250 mM sucrose were replaced with 120 mM NaCl (pH = 7.35-7.4) and incubated for at least 1.5 h to recover before recordings. Recordings were performed in a submerged chamber (Warner instruments RC-27L), supplied with continuously bubbled ACSF with a solution exchange of 3-5 ml per min at room temperature (22-24°C). An upright microscope (Olympus, BX61WI) was used for the positioning of slices to have access to the CA1 region of the hippocampus for recordings and electrode placement. Electrodes were made from glass capillaries (Hilgenberg) using the micropipette puller Sutter P-1000 (Sutter Instruments). Stimulating (1-1.5 MΩ) and recording (1.5-2.5 MΩ) electrodes filled with ACSF were placed in *stratum radiatum* of the CA1 region, and field excitatory postsynaptic potentials (fEPSPs) were recorded. For basal stimulation 0.2 ms pulses were delivered at 0.05 Hz at the stimulation intensity which induced approximately 30-50% of the maximal responses. After 10 min of stable baseline recordings stimulus response curves were made as a measure of basal excitatory synaptic transmission. Stimulation intensity was increased by 10 µA steps until the maximal fEPSP was obtained. The slopes of fEPSPs and amplitudes of presynaptic volleys were plotted vs amplitudes of presynaptic volleys and increasing stimulation intensities respectively as a measure of basal synaptic function and presynaptic excitability. The data were recorded at a sampling rate of 10 kHz, low-pass filtered at 3 kHz and analyzed using PatchMaster software and EPC9 amplifier (Heka Elektronics). SigmaPlot (Systat) software was used for data analyses and presentation.

### Generation of lysates for protein quantification by immunoblotting

All steps were performed at 4°C in the presence of protease inhibitors (complete™, Mini protease inhibitor cocktail, Merck) and phosphatase inhibitors (phosphatase inhibitor cocktail 2 and 3, Merck). Brains from 18 month-old AD/CALM^lox^ and AD/CALM^E-ko^ female littermate mice were homogenized in ice-cold RIPA buffer (150 mM NaCl, 1% NP-40, 0.5% sodium deoxycholate, 0.1% SDS, 50 mM Tris-HCl) using 20 strokes at 900 rpm. The lysates were incubated under gentle rotation for 15 min at 4°C to ensure complete lysis, prior to centrifugation at 900 x g for 10 min for removing large cellular debris. Protein concentration was measured by BCA assay. Equal protein amounts were diluted in Laemmli sample buffer, resolved by SDS-PAGE and analyzed by immunoblotting.

### Preparation of neuronal cell cultures and transfection

Cortico-hippocampal neurons or hippocampal neurons were isolated from neonatal mouse brains [postnatal days (p) 0-3] and were prepared in sparse or mass culture, respectively. Briefly, cortices and hippocampi or hippocampi alone (pooled from several genotypically identical littermates) were rapidly dissected under a binocular microscope, placed into ice-cold HEPES-buffered Hank’s balanced salt solution (HBSS, Thermo Fisher Scientific) containing 20% fetal bovine serum (FBS), and cut with a scalpel into ca. 1 mm^3^ sized pieces. The tissue pieces were washed first with HBSS containing 20% FBS and then with HBSS only and afterwards were digested for 15 min in digestion buffer (137 mM NaCl, 5 mM KCl, 7 mM Na_2_HPO_4_, 25 mM HEPES, 1 mg/ml trypsin, 1500 units DNase, pH 7.2) at 37 °C, followed by another washing step with HBSS and gentle trituration in dissociation buffer (HBSS containing 12 mM MgSO_4_, 1500 units DNase, pH 7.2). 100,000 (for each well of a 6-well plate) or 50,000 hippocampal neurons (for each well of a 12-well plate) were plated as 40 µl or 20 µl drops, respectively, per poly-L-lysine coated coverslip. 2 ml (for each well of a 6-well plate) or 1 ml (for each well of a 12-well plate) of plating medium [basic medium (MEM medium) supplemented with 0.5% glucose, 0.02% NaHCO_3_, 0.01% transferrin, 10% FBS, 2 mM L-glutamine, 25 µg/ml insulin and 1% penicillin/streptomycin] was added 1 h after plating. For biotinylation experiments, 1,200,000 cortico-hippocampal neurons (for each well of a 6-well plate) were plated as sparse culture directly in 2 ml of plating medium. After one day in vitro (DIV1), half of the plating medium was replaced by growth medium (basic medium containing 5% FBS, 0.5 mM L-glutamine, 1x B27 supplement, 1% penicillin/streptomycin). On DIV2, 1 ml (for each well of a 6-well plate) or 500 µl (for each well of a 12-well plate) of growth medium were added. 2 µM AraC were used during medium renewal to limit glial proliferation. Neurons were maintained at 37°C in a 5% CO_2_ humidified incubator until DIV12-14 for cortico-hippocampal neurons or DIV14-16 for hippocampal neurons. To initiate gene deletion, primary neurons isolated from iCAG-Cre-positive mice (CALM^cag-ko^) were treated with 0.3 µM (Z)-4-hydroxytamoxifen (Sigma-Aldrich) immediately after plating. An equal concentration of tamoxifen (0.3 µM) was used during medium renewal on DIV1 and DIV2. Neurons derived from floxed littermates that were Cre negative (CALM^lox^) were used as controls and were treated with equal amounts of (Z)-4-hydroxytamoxifen. For transient expression of VAMP8-pHluorin (kind gift of Dr. Rytis Prekeris), neurons were transfected on DIV7-8 by calcium phosphate (ProFection® Mammalian Transfection System, Promega) using 2 µg of plasmid DNA, 250 mM CaCl_2_ and nuclease-free water (for each well of a 6-well plate) mixed with equal amounts of 2X HEPES-buffered saline (2x HBS, 100 µl). The mix was incubated at room temperature (RT) for 20 min to allow precipitate formation while neurons were starved in osmolarity-adjusted NBA (Neurobasal™ Medium, Thermo Fisher Scientific) at 37°C, 5% CO_2_ for the same time. Precipitates were added to neurons and incubated for 20 min at 37°C, 5% CO_2_. Finally, neurons were washed two times with osmolarity-adjusted HBSS (Hank’s Balanced Salt Solution, Thermo Fisher Scientific) and transferred back to their conditioned medium, which had been previously saved. Live imaging was conducted at 7-8 days post-transfection (DIV 14-16).

### Surface biotinylation of cortico-hippocampal cultures

To biotinylate surface proteins, after washing once with ice-cold PBS^2+^ (137 mM NaCl, 2.7 mM KCl, 10 mM Na_2_HPO_4_, 1.8 mM KH_2_PO_4_, 1 mM CaCl_2_, 0.5 mM MgCl_2_, pH 7.4), cortico-hippocampal neurons at DIV12-14 in 6-well plates were incubated in the dark with 0.5 mg/ml of the non-membrane-permeant, cleavable biotin derivative sulfo-NHS-SS-biotin for 20 min under gentle shaking at 4°C. After biotinylation, neurons were washed twice with ice-cold quenching solution (50 mM glycine in PBS^2+^) for 5 min to remove excess biotin. Then, neurons were rinsed once with ice-cold PBS^2+^ and immediately solubilized in RIPA buffer supplemented with protease inhibitors (cOmplete™, Mini protease inhibitor cocktail, Merck) and phosphatase inhibitors (phosphatase inhibitor cocktail 2 and 3, Merck). Lysates were cleared by centrifugation at 13,000 x g for 10 min at 4°C, supernatants were collected, and protein concentration was determined by BCA assay. A portion of the supernatant was used as input. To precipitate the biotinylated proteins, equal amounts of total protein were incubated with streptavidin-agarose beads (Thermo Fischer Scientific) at 4°C with rotation. After 2 h beads were extensively washed with RIPA buffer (150 mM NaCl, 1% NP-40, 0.5% sodium deoxycholate, 0.1% SDS, 50 mM Tris-HCl), and surface biotinylated proteins were eluted by boiling for 10 min in Laemmli sample buffer, resolved by SDS-PAGE and immunoblotted using appropriate antibodies. Equal amounts of total cell lysates were analyzed for total protein levels. Ratiometric quantification of signal intensities was performed with the Image studio software package of the Odyssey Fc Imaging system (LI-COR Biosciences).

### Aβ uptake assay and immunostaining of hippocampal neurons in culture

HiLyte Fluor™ 488-labeled Aβ_42_ (Anaspec Inc., AS-60479-01) was reconstituted by adding 50 μl 1% NH_4_OH to 0.1 mg peptide. This peptide solution was diluted to 1 mg/ml with 1x PBS, aliquoted and stored at -20°C. To investigate the internalization of Aβ_42_, hippocampal neurons were incubated with the labeled Aβ_42_ at 500 nM for 6 h. Neurons were then washed twice with PBS and fixed with 4% (w/v) PFA and 4% sucrose in 1x PBS for 10 min at room temperature (RT). Neurons were then permeabilized for 5 min using 0.1% Triton X-100, blocked with 2% BSA in 1x PBS and incubated with the indicated primary antibodies diluted in 2% BSA in 1x PBS overnight at 4°C, followed by washes and then a 1-hour incubation at room temperature with the appropriate secondary antibodies diluted 1:400 in 1x PBS. Neuronal nuclei were visualized with DAPI (1 μg/ml in 1× PBS), and coverslips were mounted using Immu-Mount (Thermo Fisher Scientific). Aβ uptake was analyzed using a Zeiss LSM 710 laser scanning confocal microscope with a 63× oil-immersion objective.

### Plasmid cloning for the generation of genetically modified *C. elegans* strains

Expression plasmids were generated using standard cloning techniques. The UPNp promotor sequence was amplified from genomic DNA of the OH15263 strain. The human PICALM sequence was amplified from the plasmid CALM-pmCherry-N1 (kind gift of David Perrais, University of Bordeaux). All fragments were assembled into an mNeongreen containing expression plasmid using restriction digestion and ligation, resulting in the following nematode expression plasmid: pJK124 (UPNp::PICALM-mNG), used in the generation of the strain JKM184 (Supplementary Table 3).

### Generation of genetically modified *C. elegans* strains

The transgenic strain JKM184 (see Supplementary Table 3) was generated via microinjection. Day 4 old wild-type nematodes were injected with the following injection mix: jksEx112 (15 ng/µl UPNp::PICALM-mNG [pJK124] + 50 ng/µl coel::RFP + 100 ng/µl 100 bp-ladder). Transgenic offspring carrying extra-chromosomal arrays were isolated and cultured.

### Nematode maintenance

Nematodes were cultured on nematode growth medium (NGM) agar plates at 20°C under standard conditions (Wood, 1988) unless stated otherwise. All age statements given in this publication consider the day of hatching as day 1.

### Fluorescence lifetime imaging

Seven-day-old nematodes expressing mScarlet or mScarlet-Aβ ± CALM (JKM2, JKM47, JKM197, JKM221) were anesthetized with 250 mM NaN_3_ for up to 30 min and mounted on a 2.5% agarose-pad before performing fluorescence lifetime measurements on a confocal microscope. Measurements were carried out using a ‘Leica SP8′ confocal microscope, equipped with a ‘HC PL APO 63 CS2 × 1.4′ objective. Zoom was applied to 1.5-fold. Images were recorded by time correlated single photon counting (TSCPC) using ‘FALCON′ (Leica). A pulsed supercontinuum laser was used for excitation, emitting pulses at 80 MHz. An acousto-optic beam splitter was used to set excitation light to 561 nm and to measure emission between 575– 625 nm. Photon count rates were kept below 1% of the laser repetition rate to prevent photon pileup. Images were acquired until approx. 4000 photons per pixel were recorded. Images were processed with ‘FlimFit 5.1.1’ software (Warren et al., 2013) and fitted assuming mono-exponential decays.

### Chemotaxis assay

Nematodes were age-synchronized by picking L4-stage larvae one day before the assay. 25-100 animals were employed per cohort and strain. Four day-old nematodes were washed off the plates with M9 buffer (3 g KH_2_PO_4_, 6 g Na_2_HPO_4_, 5 g NaCl, 1 ml 1 M MgSO_4_ ad 1 l H_2_O)(Conte et al., 2015) and then washed three times for 5 min with M9 buffer. Nematodes were left to starve for 30 min in M9 buffer after the final wash. Non-seeded 6 cm NGM plates were labeled with four quadrants and a 1 cm circle in the middle. 1% (v/v) benzaldehyde was mixed with 500 mM NaN_3_ (ratio 1:1), and 2 µl were applied on two opposing quadrants. In the remaining quadrants 2 µl of solvent control (water) mixed with 500 mM NaN_3_ were spotted. Washed nematodes were placed in the middle circle and allowed to move according to their preference for 1 h at RT. Afterwards, the chemotactic index was calculated as the difference between the fraction of nematodes on the sample quadrants and on control quadrants, divided by the total number of nematodes counted.

### Aβ42 expression and purification for Thioflavin T assay

Aβ42 peptide was overexpressed and purified as described previously (Ayala Mariscal et al., 2022; Hoarau et al., 2016). In short, 50 ng of pET-M-Aβ42 plasmid were transformed into *E. coli* BL21-CodonPlus (DE3)-RIPL cells (Agilent, #230280). A starter culture was prepared by inoculating 30 ml of LB media supplemented with Ampicillin (100 μg/ml) with 5 transformants followed by incubation at 37°C and shaking at 180 rpm overnight. The next morning, 1 l of LB-Ampicillin media was inoculated with 10 ml of the starter culture and incubated at 37°C and shaking at 180 rpm. When OD600 reached 0.6, protein expression was induced by adding 0.5 mM IPTG. After 4 additional h of incubation, the cultures were centrifuged at 20.000 g (Sorvall Evolution RC, Thermo Scientific) for 20 min at 4°C. The resulting pellets were resuspended in 100 ml lysis buffer (50 mM Tris-HCl pH 8.0, 100 mM NaCl, 1% NP40, 2 mM PMSF and 1 tablet/50 ml of cOmplete EDTA-free protease inhibitor cocktail (Roche)) and subjected to 4 cycles of cell disruption at 16000 psi (LM10 Microfluidizer, Microfluidics). The insoluble fraction containing the Aβ_42_ peptide was recovered by centrifugation at 12.000×g (Sorvall Evolution RC, Thermo Scientific) for 30 min at 4°C. To clean the peptide from cellular debris, the insoluble fraction was resuspended in cleaning buffer (lysis buffer supplemented with 300 µg/ml of lysozyme and 2 mM EDTA) and incubated for 1 h at RT with gentle rotation. The suspension was next centrifuged at 12.000×g for 10 min at 4°C, and the pellet was washed twice by resuspension in 35 ml wash buffer 1 (PBS pH 8.0, 0,5% Triton X-100), followed by incubation for 10 min at RT with gentle rotation and centrifugation at 12.000×g for 10 min. Next, two wash steps were performed in 35 ml wash buffer 2 (PBS pH 8.0), followed by ten cycles of 10 s sonication at 50% amplification (HD 2070, Bandelin Sonoplus). The suspension was divided into four equal volumes and centrifuged at 12.000×g for 10 min at 4°C. To solubilize the peptide, each pellet was resuspended in 35 ml of 50 mM NaOH and incubated for 2 h at RT with gentle rotation, followed by centrifugation at 12.000×g for 10 min at 4°C. The supernatant was recovered, and its pH was carefully adjusted to 7.0 by slow addition of 0.5 M HCl. The suspension was centrifuged at 12.000×g for 10 min at 4 °C, and the supernatant containing the soluble Aβ_42_ was recovered. The remaining pellet was subjected to another round of NaOH solubilization and pH neutralization. The recovered supernatants were pooled and filtered through a 30 kDa MWCO filter (Amicon Ultra-15 Centrifugal filters, Millipore). The flow-through was concentrated to around 4–5 ml in a 3 kDa MWCO filter (Amicon Ultra-15 Centrifugal filters, Millipore), and 1 ml aliquots were prepared and flash-frozen in liquid nitrogen for storage at −80 °C.

### Expression and purification of His-tagged proteins for Thioflavin T assay

His-CALM-ANTH and His-Δ51-Gadkin proteins were expressed in *E. coli* (BL21) at 16°C for 16-18 h. The pelleted cells were lysed, and cell debris was removed by centrifugation. For purification, the cleared supernatants were incubated with His-Select Nickel Affinity Gel (Sigma-Aldrich). After multiple washing steps, the proteins were eluted according to the manufacturer’s instructions.

### Thioflavin T assay to monitor Aβ42 aggregation

Prior to the Thioflavin T assay, proteins were thawed on ice and transferred to centrifugation tubes (Microfuge Tube Polypropylene, Beckman Coulter). Ultracentrifugation (Optima MAX Ultracentrifuge, Beckman) was carried out at 120.000 g for 40 min at 4°C. Protein supernatants (around 95% of the initial volume) were transferred to low-binding tubes (Sorenson), and the molar concentration was determined via Bradford assay. Subsequently, Thioflavin T (ThT) assays were performed as described in (Ayala Mariscal et al., 2022). After ultracentrifugation, Aβ42 was loaded onto a size-exclusion column (Superdex 75 10/300) previously equilibrated with 50 mM NaOH. The peptide, which elutes at around 13 ml, was recovered, concentrated and its molar concentration was determined via Bradford assay. 160 µl samples were prepared containing 30 mM HEPES pH 7.4, 150 mM NaCl, 5 mM MgCl_2_, 10 µM ThT, 5 µM Aβ_42_, and 5 or 1 µM His-CALM-ANTH or His-Δ51-Gadkin. In total, 50 µl aliquots of each sample were transferred into three different wells of a 384-well non-binding black microplate (Corning, #3575). The microplate was sealed with a transparent coverslip and ThT fluorescence was measured every 10 min over a period of 40 h using a plate reader (Infinite F200 PRO, Tecan) set at 37°C. Excitation and emission wavelengths were 440 nm and 490 nm, respectively, and a 10 s and 2 mm shaking cycle was done before each measurement. ThT fluorescence was plotted as a function of time, and all values were normalized to the sample of Aβ42 in the absence of CALM or Gadkin.

### Blot analysis of soluble fractions

The samples of Aβ42 incubated in the presence of CALM-ANTH or Gadkin from the previously described ThT assay were recovered after 40 h. Soluble and aggregated Aβ42 fractions were separated by centrifugation at 120.000 g for 40 min at 4°C. Supernatants were recovered and blotted in triplicates on a PVDF membrane using a dot-blot device. A sample of fresh monomeric Aβ42 was used as control. Blotted membranes were blocked for 1 h with TBST containing 5% BSA. Primary and secondary antibodies were mouse anti-Aβ (clone 6E10), and anti-mouse antibodies coupled to HRP, respectively. Assay signals were developed using Pierce ELC (Thermo Fisher) and detected on an OdysseyFC imaging system (Image Studio TM software 4.0; LI-COR Biosciences). Image analysis was performed using Fiji/ImageJ (Schindelin et al., 2012).

### pHluorin imaging of living hippocampal neurons

Hippocampal neurons were imaged at RT in N-Tris(hydroxy-methyl)-methyl-2-aminoethane-sulphonic acid (TES)-based physiological imaging buffer (170 mM NaCl, 3.5 mM KCl, 0.4 mM KH_2_PO_4_, 20mM TES, 5 mM NaHCO_3_, 5 mM glucose, 1.2 mM Na_2_SO_4_, 1.2mM MgCl_2_, 1.3mM CaCl_2_, pH 7.4) with an epifluorescence microscope (Nikon Eclipse Ti with eGFP filter set F36-526 and an sCMOS camera (Neo, Andor)) equipped with a 40x oil-immersion objective and operated by open-source ImageJ-based MicroManager 4.11 software. In initial optimization experiments, the imaging parameters were established to ensure minimal photobleaching of fluorescence. Fluorescence of pHluorin–tagged proteins was stable during the imaging experiments (time, >5 min). Images were acquired every 2 s with 100 ms excitation at 488 nm. To measure the steady state surface to total ratios, the surface-localized pHluorins (FS) were first quenched by replacing the extracellular TES solution by MES [2-(N-morpholino) ethanesulfonic acid] (pH 5.5). To measure the total fluorescence of overexpressed pHluorins (FT), 50 mM NaCl were replaced by NH_4_Cl. Regions for analysis were selected manually, and quantitative analysis was performed using ImageJ (NIH). The surface to total ratio was calculated as (baseline - FS)/(FT - FS).

### Purification of synaptosomes and TMT-based mass spectrometry

All steps were performed at 4°C in the presence of protease inhibitors (cOmpleteTM, EDTA-free protease inhibitor cocktail tablet, Sigma-Aldrich). Mouse brains from 3 AD/CALM^lox^ and 3 AD/CALM^E-ko^ 14-15 month-old female littermate mice were homogenized in ice-cold sucrose buffer (320 mM sucrose, 20 mM HEPES, 5 mM EDTA, pH 7.4) with 12 strokes at 900 rpm. Samples were then centrifuged at 900 × g for 10 min. The supernatant (S1) was collected; the resulting pellet (P1) contains large cell fragments and nuclei. S1 was then centrifuged at 10,000 × g for 15 min. The supernatant (S2) was removed, whereas the pellet (P2), containing synaptosomes, was carefully resuspended in 4 ml of sucrose buffer and centrifuged at 10,000 × g for 15 min (washing step). The washed crude synaptosomal pellet (P2’) was resuspended in 1.5 ml sucrose buffer. Protein concentration was measured by BCA assay. Samples were supplemented with 50 mM Triethylammonium bicarbonate buffer (TEAB) pH 8.5, 8 M urea and benzonase, and homogenized for 10 min in a Bioruptor (30 s on /30 s off). Subsequently, disulfide bonds were reduced with 5 mM Tris(2-carboxyethyl)phosphine (TCEP) and alkylated with 40 mM chloroacetamide (CAA)for 1 h at 37°C in the dark. Proteins were first digested using LysC (1:200, wt:wt, enzyme:protein ratio) for 2 h at 37°C. The urea concentration was diluted to final 2 M using 50 mM TEAB, and samples were further digested with trypsin (1:100, wt:wt, enzyme:protein ratio) for 16 h at 37°C. The digestion was stopped by addition of 10% formic acid (FA) to a final concentration of 1%, and peptides were desalted using C18 SepPak. For each sample, 120 µg of peptides were resuspended in 25 µl of 50 mM TEAB pH 8.5. Then 320 µg TMT6-plex reagent (Thermo Scientific) in 16 µl acetonitrile (ACN) were added to each sample. Labeling was performed for 1 h at RT. The reaction was quenched with 46 mM Tris-HCl pH 8.0 for 15 min at RT. Labeling efficiency of >98% was confirmed, and samples were mixed at 1:1 ratio according to a mixing control adjusted to equal median TMT reporter ion intensities. The pooled sample was dried briefly to remove excess ACN and desalted using C18 SepPak. 200 µg peptides were resuspended in 10 mM NH_4_OH in water and injected into a Gemini C18 column (3 µm, 100 × 1 mm, phenomenex) installed on an Agilent 1260 Infinity II UPLC. At a flow rate of 0.1 ml/min, peptides were separated over 85 min using increasing concentrations of 10 mM NH_4_OH in ACN as an eluent. Finally, all fractions were dried and stored at - 20°C until measurement. Dried peptide fractions were resuspended in 1% ACN and 0.05% trifluoroacetic acid (TFA) and neighboring fractions merged to yield 12 separate injections. For each run, 1 µg of peptides was injected into a Dionex Ultimate3000 HPLC system connected to a PepMap C18 trap column (75 µm × 50 mm, 3 µm particle size, 100 Å pore size, Thermo Fisher Scientific), followed by an in-house packed C18 analytical column (Poroshell 120 EC-C18, 2.7 µm, Agilent Technologies). Peptides were separated at a flow rate of 250 nl/min using a 120-min gradient with increasing ACN concentration and analyzed on an Orbitrap Fusion Lumos mass spectrometer using Instrument Control Software version 4.0. MS1 scans were acquired in the Orbitrap at a resolution of 120,000 over a scan range of m/z 400–1,600, with a 100% AGC target and 246 ms maximum injection time. MS2 scans were acquired in the Orbitrap with the following parameters: first mass 110 m/z, 250% AGC target, 86 ms maximum injection time, isolation window 0.7 m/z, normalized collision energy (NCE) 38%, and resolution 50,000. Previously isolated precursors were dynamically excluded for 60 s. Only precursors with charge states +2 to +6 were selected for MS2. Data was acquired using 2 s per cycle. Raw files from high-pH fractions of TMT6-plex labeled synaptosome-derived peptides were analyzed using FragPipe v23.1 with MSFragger v4.3 and IonQuant v1.11.11. Spectra were searched against the murine proteome retrieved from UniProt (downloaded 2025-09-02, one protein per gene). Precursor and fragment ion mass tolerances were set to ± 10 ppm and ± 20 ppm, respectively, after mass recalibration. Trypsin was specified as the protease with up to two missed cleavages allowed. Protein N-terminal methionine clipping was enabled. Variable modifications included methionine oxidation (+15.9949 Da), protein N-terminal acetylation (+42.0106 Da), TMT labeling (+229.16293 Da) on peptide N-termini and serine residues. Fixed modifications included carbamidomethylation of cysteines (+57.02146 Da) and TMT labeling of lysine residues. Peptide length was restricted to 7–50 amino acids and peptide mass to 200–5,000 Da. All other parameters were kept at default settings from the TMT-workflow. TMT reporter ion intensities were extracted with IonQuant using a ± 20 ppm tolerance, no normalization, MS2-based quantification, best PSM selection, allowance of overlabeling, and outlier removal. False discovery rate (FDR) was controlled at 1% using Percolator and MSBooster. For quantitative analysis, gene TMT report tables from FragPipe were imported into R. Intensities were log2-transformed and median-normalized. Fold changes and p-values were calculated using a moderated Bayesian t-test implemented in the limma R package.

### Statistics and reproducibility

Data are depicted as individual data points and mean±SEM and represent values obtained from individual animals respectively independent experiments as indicated in the figure legends and in the Source Data file. Statistical evaluation of differences between groups was based on data from at least three animals or three independent experiments. Data distribution was tested by Shapiro-Wilk test. Data where each control value was normalized to 100 were tested by two-tailed One sample t test. Comparisons of two groups of normally distributed data were either performed by two-tailed paired t test or by two-tailed unpaired t test with Welch’s correction. Comparisons of two groups of not normally distributed data were done by Wilcoxon matched-pairs signed rank test or by Mann Whitney test. For comparisons of more than two groups of normally distributed data, the one-way ANOVA was used combined with Holm-Sidak’s multiple comparison test. For comparisons of more than two groups of not normally distributed data the Kruskal-Wallis test was used combined with Dunn’s multiple comparison test. Electrophysiological data was analyzed by two way repeated measures ANOVA. The specific statistical tests chosen is provided within the figure legends and in the Source Data File which also states exact p values. Throughout the figures significance levels are indicated as *=p<0.05; **=p<0.01; ***=p<0.005; ****=p<0.0001. Differences that are not significant are indicated as ns. Statistical data evaluation was performed using GraphPad Prism (version 11.0.2).

## Supporting information

Supplementary Table 1 - Primers

Supplementary Table 2 - Antibodies

Supplementary Table 3 - Nematode Strains

Compilation of Uncropped Blots

## Acknowledgements

We thank Silke Zillmann for expert technical assistance. Schematics were created with BioRender.com.

## Author contribution

DAT, JK, VH and TM conceived and designed experiments. DAT, GK, CG, MS, SAM, JKS, KG, KK,

CB, DL, GC performed research. FL supervised mass spectrometry experiments conducted by HS, MW, DAT and MR. DAT, JK, VH, TM wrote the manuscript with input from all authors.

## Funding

This research was funded by grants of the Deutsche Forschungsgemeinschaft (DFG, German Research Foundation) to TM (MA4735/1-2), to JK (KI1988/7-1) and VH (HA2686/8-2) and by Forschungsinitiative Rheinland-Pfalz BioComp to TM. DAT was supported by an Alexander von Humboldt Research Fellowship for Postdoctoral Researchers and a Klaus Tschira Boost Fund. DAT is now supported by the Programma per Giovani Ricercatori “Rita Levi Montalcini” of the Italian Ministry of University and Research, by the University of Padova (STARS@UNIPD 2025 and Progetto DOR 2025/2026), and by a Career Support Grant 2025 from the International Society of Neurochemistry (ISN). Mass spectrometry experiments were supported by the European Research Council (ERC) Starting Grant (ERC-STG no. 949184) to FL and by funding via the China Scholarships Council (no. 202306310041) to MW.

## Ethics

All mouse experiments were reviewed and approved by the ethics committee of the “Landesamt für Gesundheit und Soziales” (LAGeSo) Berlin and were conducted according to the committee’s guidelines under animal experimentation permit T0243/08. At the institute, the Animal Care Officer and the LAGeSo monitored compliance with all regulations.

## Competing interests

The authors declare no competing interests.

## Supplementary Information

can be found online.

## Notes

### Competing Interest Statement

The authors have declared no competing interest.

