## Supplementary Figures for "PICALM deficiency exacerbates amyloid pathology in murine and nematode Alzheimer’s disease models"

Items contained in this file:

Supplementary Figure 1:
**Characterization of AD/CALM$^{E\text{-}ko}$ mice**

Supplementary Figure 2:
**Description and representative images of animal models**

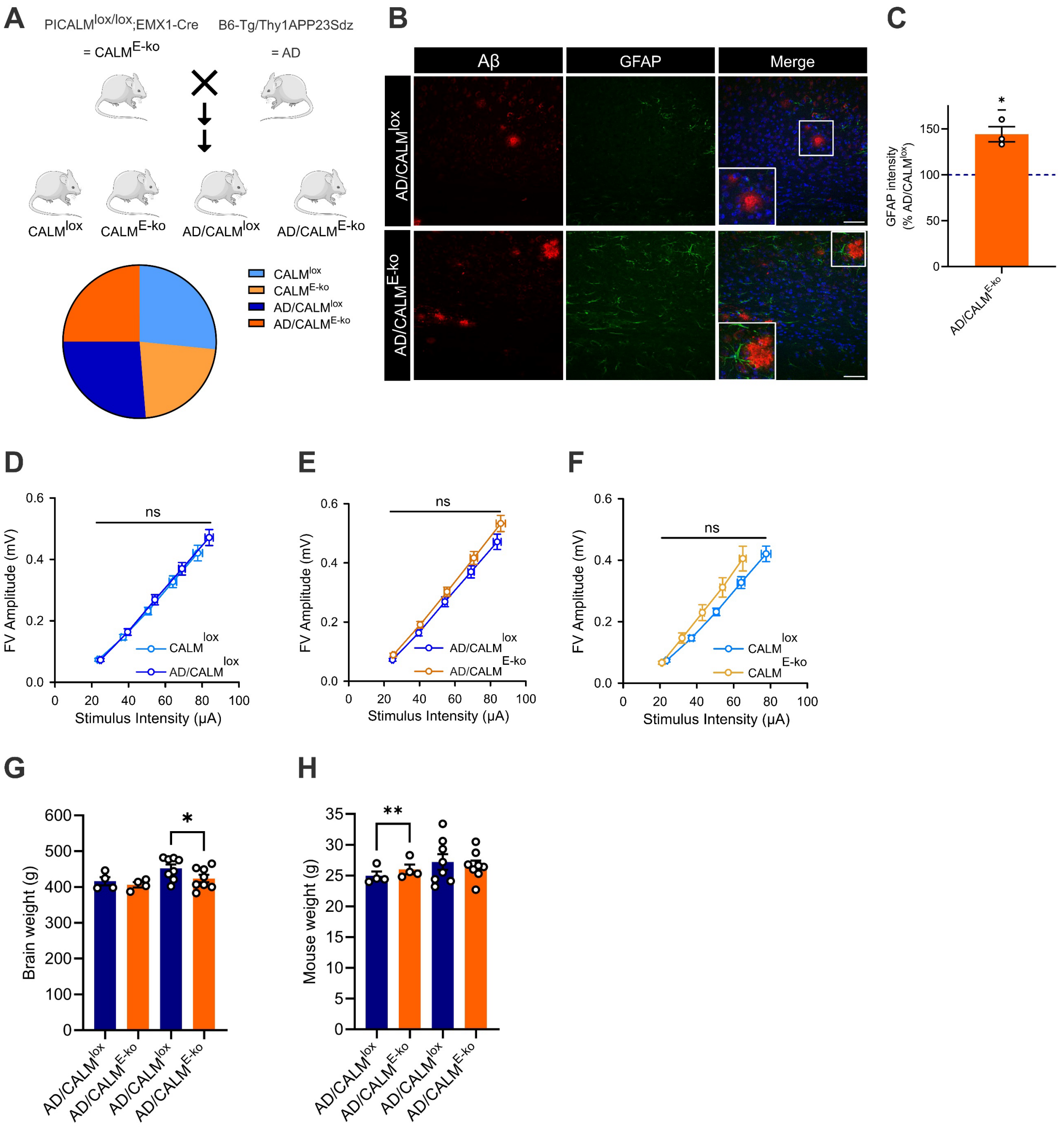


**Supplementary Fig. 1 Characterization of AD/CALM$^{E-ko}$ mice.**

**(A)** Top: Illustration of breeding scheme for AD/CALM$^{E-ko}$ mice. Bottom: Mendelian distribution of offspring. **(B-C)** Loss of CALM aggravates GFAP-positive Ab aggregates. **(B)** Representative images of hippocampal brain sections of AD/CALM$^{lox}$ and AD/CALM$^{E-ko}$ mice stained with Aβ-, GFAP- and MAP2-specific antibodies and DAPI to label nuclei. Scale bar, 50 µm. **(C)** Quantification of GFAP levels in AD/CALM$^{E-ko}$ brains normalized to AD/CALM$^{lox}$ controls. Dots represent individual animals. One sample t test. **(D-F)** CALM loss does not alter fiber volley excitability in any of the tested genotypes. **(D)** The relationship between fiber volley amplitudes and stimulus intensities reveal no significant difference in AD/CALM$^{lox}$ mice as compared to CALM$^{lox}$ mice. Two-way repeated measures ANOVA, AD/CALM$^{lox}$: N=8 animals/n=21 slices; CALM$^{lox}$: N=9 animals/n=21 slices. **(E)** The relationship between fiber volley amplitudes and stimulus intensities reveal no significant difference in AD/CALM$^{E-ko}$ mice as compared to AD/CALM$^{lox}$ mice. Two-way repeated measures ANOVA, AD/CALM$^{E-ko}$: N=9 animals /n=24 slices; AD/CALM$^{lox}$: N=8 animals/n=21 slices. **(F)** The relationship between fiber volley amplitudes and stimulus intensities reveal no significant difference between CALM$^{lox}$ and CALM$^{E-ko}$ mice. Two-way repeated measures ANOVA, CALM$^{lox}$: N=9 animals/n=21 slices; CALM$^{E-ko}$: N=6 animals/n=16 slices. **(G)** Decreased brain weight in 18-month-old female AD/CALM$^{E-ko}$ mice as compared to AD/CALM$^{lox}$ mice (two right columns), while there is no difference yet at 12 months (two left columns). Two-tailed paired t-tests within age groups. Circles denote individual animals, N=4-8. **(H)** No decrease in body weight in 12- (two left columns) and 18- (two right columns) month-old female AD/CALM$^{E-ko}$ mice as compared to AD/CALM$^{lox}$ mice. The body weight of 12-month-old female AD/CALM$^{E-ko}$ mice as compared to AD/CALM$^{lox}$ mice is even slightly increased. Two-tailed paired t tests within age groups. Circles denote individual animals, N=4-8. For all graphs data are depicted as mean±SEM. $*=p<0.05$; $**=p<0.01$; ns, non significant.

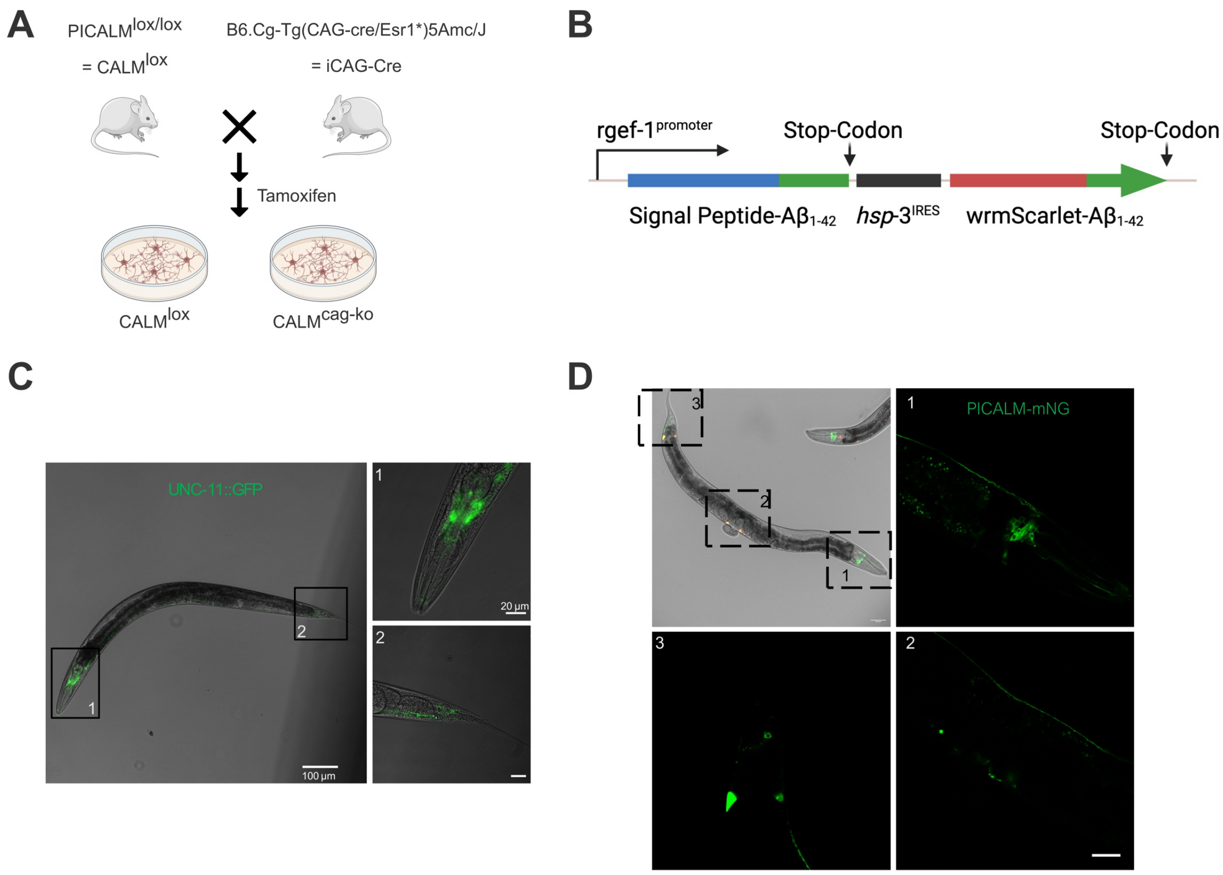


**Supplementary Fig. 2. Description and representative images of animal models.**
**(A)** Illustration of breeding scheme for inducible CALM ko mouse line. The ko is induced in cell cultures by tamoxifen addition. **(B)** Illustration of *C.elegans* Alzheimer´s disease model showing the organization of the operon driving the overexpression of secreted human $A\beta_{42}$ and the internal ribosomal entry site (IRES (derived from hsp-3))-dependent sub-stoichiometric expression of mScarlet-$A\beta_{42}$ under the pan-neuronal promoter rgef-1p (strain JKM2). Modified from Gallrein et al., 2021. **(C-D)** Representative confocal images of used *C.elegans* strains. **(C)** The strain JKM45 expresses the *C.elegans* version of CALM tagged with GFP (*unc-11::gfp)* pan-neuronally under the rgef-1 promotor (JKM45 – *jksIs45* [*rgef-1p*::*unc-11*$^{isoform\ a}$::GFP]). **(D)** The strain JKM184 expresses the human CALM protein tagged with mNeongreen pan-neuronally under the UPN synthetic ultra-pan-neuronal promotor and also contains a fluorescent selection marker (RFP), which labels coelomocytes (JKM184 – *jksEx112* [UPNp::PICALM-mNeon-Green + coel::RFP]). Confocal images represent merges of brightfield and green fluorescence channel at 100x magnification; insets are 630x magnifications of indicated nematode body regions. Scale bar, 100 µm. Scale bar in insets, 20 µm.
