## Supplementary Table 1 - Primers for "PICALM deficiency exacerbates amyloid pathology in murine and nematode Alzheimer’s disease models"

**Supplementary Table 1. Genotyping Primers**

| Mouse line | Primer name | Sequence |
|---|---|---|
| PICALM LOX | TM435 | GTGGGATGAATGGTTGGGCTTC |
| | TM436 | CCATGTAGGGTCTAAAGATC |
| EMX Cre and iCAG Cre | TM63 | CCGGGCTGCCACGACCAA |
| | TM64 | GGCGCGGCAACACCATTTTT |
| APP23 | APP23_fwd | GAATTCCGACATGACTCAGG |
| | APP23_rev | GTTCTGCTGCTGCATCTTGGACA |
