## Supplementary Table 2 - Antibodies for "PICALM deficiency exacerbates amyloid pathology in murine and nematode Alzheimer’s disease models"

| Antigen | Species | Dilution WB | Dilution IF | Supplier |
|---|---|---|---|---|
| APP (6E10) | Rabbit | | 1:200 | Novus Biologicals, NBP2-62566 |
| APP (Y188) | Rabbit | 1:1000 | 1:500 | Abcam, ab32136 |
| BACE1 | Rabbit | 1:1000 | | Cell Signaling, #5606 |
| $\beta$-Actin | Mouse | 1:5000 | | Sigma-Aldrich, A5441 |
| CALM | Rabbit | 1:500 | | Novus Biologicals, NBP1-86658 |
| GFAP | Mouse | 1:000 | 1:400 | Synaptic Systems, 173211 |
| HSP70 | Mouse | 1:2000 | | Invitrogen, MA3-006 |
| LRP1 | Rabbit | 1:1000 | | Abcam, ab92544 |
| MAP2 | Guinea pig | | 1:500 | Synaptic System, 188004 |
| Neurofilament M | Mouse | | 1:500 | Synaptic System, 171211 |
| N-cadherin | Mouse | 1:500 | | BD Biosciences, 610920 |
| Nicastrin | Rabbit | 1:500 | | Cell Signaling, #3632 |
| Presenilin 2 | Rabbit | 1:200 | | Abcam, ab51249 |
| Syntaxin7 | Sheep | 1:500 | | Novus Biologicals, AF5478 |
| VAMP8 | Rabbit | 1:500 | | Abcam, ab76021 |
| IRDye 680RD donkey anti-mouse IgG secondary antibody | Mouse | 1:10000 | | LI-COR Biosciences, 926-68072 |
| IRDye 800CW donkey anti-rabbit IgG secondary antibody | Rabbit | 1:10000 | | LI-COR Biosciences, 926-32213 |
| Goat Anti-Guinea Pig IgG (H+L) Highly Cross-Adsorbed Secondary Antibody, Alexa Fluor 647 | Guinea pig | | 1:500 | Thermo Fisher Scientific, A-21450 |
| Goat Anti-Mouse IgG (H+L) Cross-Adsorbed Secondary Antibody, Alexa Fluor 488 | Mouse | | 1:500 | Thermo Fisher Scientific, A-11001 |
| Goat anti-Rabbit IgG (H+L) Cross-Adsorbed Secondary Antibody, Alexa Fluor™ 568 | Rabbit | | 1:500 | Thermo Fisher Scientific, A-11011 |

WB: Western Blotting; IF: Immunofluorescence
