## Supplementary Table 3 - Nematode Strains for "PICALM deficiency exacerbates amyloid pathology in murine and nematode Alzheimer’s disease models"

**Supplementary Table 3.**
**Overview of the genetically modified *C.elegans* strains used in this study**

| Strain name | Genetic modification | Short name |
|---|---|---|
| JKM2 | pan-neuronal overexpression of secreted human Aβ42 and sub-stoichiometric expression of mScarlet-Aβ42 under the rgef-1 promotor (*jksIs2* [*rgef-1p::Signalpeptide-Aβ42::hsp-3(IRES)::wrmScarlet-A*$\beta_{42}$*::unc-54(3'UTR) + rps-0p::HygroR*]; generated by (Gallrein et al., 2021)) | nAβ |
| JKM45 | pan-neuronal overexpression of GFP-tagged unc-11 under the rgef-1 promotor (*jksIs45* [*rgef-1p::unc-11isoform a::GFP*]; generated in this study) | unc-11$^{OE}$ |
| JKM184 | pan-neuronal overexpression of mNeongreen-tagged human CALM under the synthetic ultra-pan-neuronal (UPN) promotor and expression of fluorescent selection marker RFP in coelomocytes (*jksEx112* [*UPNp::PICALM-mNeonGreen* + *coel::RFP*]; generated in this study) | nCALM$^{OE}$ |
| CB47 | *unc-11* knockout (generated by (Brenner, 1974)) | Δunc-11 |
| JKM47 | Genetic cross of JKM2 and CB47 (generated in this study) | Δunc-11 x nAβ |
| JKM197 | Genetic cross of JKM2 and JKM184 (generated in this study) | nCALM$^{OE}$ x nAβ |
| JKM221 | Genetic cross of JKM2 and JKM45 (generated in this study) | unc-11$^{OE}$ x nAβ |

**Associated literature**:

Brenner, S., 1974. *The genetics of Caenorhabditis elegans*. Genetics. 77, 71-94.
Gallrein, C., et al., 2021. *Novel amyloid-beta pathology C. elegans model reveals distinct neurons as seeds of pathogenicity*. Prog Neurobiol. 198, 101907.
