## Supplementary figures and images for "PICALM deficiency exacerbates amyloid pathology in murine and nematode Alzheimer’s disease models"

### Compilation of Uncropped Blots

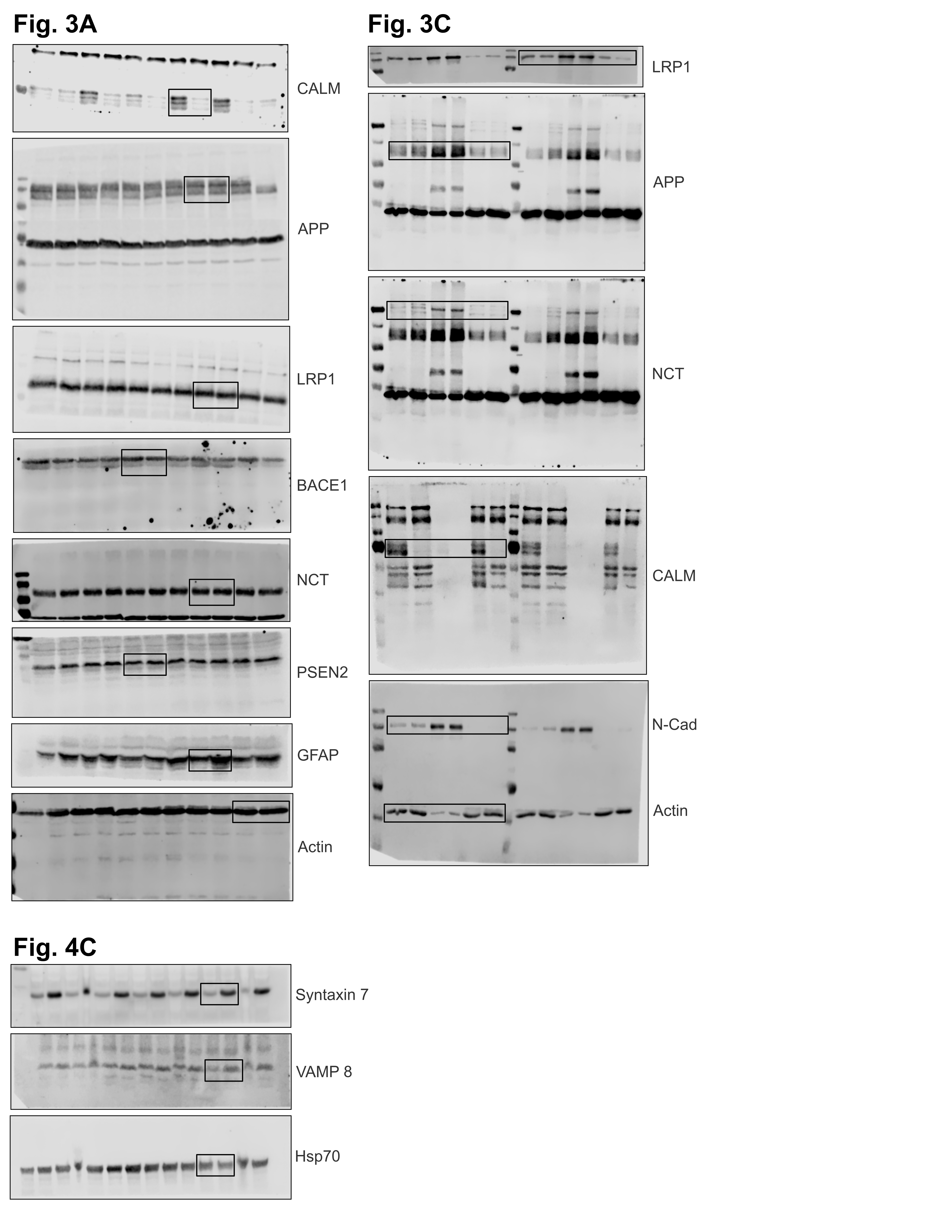
